# Mechanism of RPA phosphocode priming and tuning by Cdk1/Wee1 signaling circuit

**DOI:** 10.1101/2025.01.16.633180

**Authors:** Poonam Roshan, Vikas Kaushik, Ayush Mistry, Abhinav Vayyeti, Aidan Antony, Riley Luebbers, Jaigeeth Deveryshetty, Edwin Antony, Sofia Origanti

## Abstract

Replication protein A (RPA) is a heterotrimeric single-strand DNA binding protein that is integral to DNA metabolism. Segregation of RPA functions in response to DNA damage is fine-tuned by hyperphosphorylation of the RPA32 subunit that is dependent on Cyclin-dependent kinase (Cdk)-mediated priming phosphorylation at the Ser-23 and Ser-29 sites. However, the mechanism of priming-driven hyperphosphorylation of RPA remains unresolved. Furthermore, the modulation of cell cycle progression by the RPA-Cdk axis is not clearly understood. Here, we uncover that the RPA70 subunit is also phosphorylated by Cdk1 at Thr-191. This modification is crucial for the G2 to M phase transition. This function is enacted through reciprocal regulation of Cdk1 activity through a feedback circuit espoused by stabilization of Wee1 kinase. The Thr-191 phosphosite on RPA70 is also crucial for priming hyperphosphorylation of RPA32 in response to DNA damage. Structurally, phosphorylation by Cdk1 primes RPA by reconfiguring the domains to release the N-terminus of RPA32 and the two protein-interaction domains that markedly enhances the efficiency of multisite phosphorylation by other kinases. Our findings establish a unique phosphocode-dependent feedback mechanism between RPA and RPA-regulating kinases that is fine-tuned to enact bipartite functions in cell cycle progression and DNA damage response.

## INTRODUCTION

Replication Protein A (RPA) is a single-strand DNA (ssDNA) binding protein that functions in almost all aspects of DNA metabolism including repair, replication, and recombination^1,2^. To serve the myriad functions on DNA, RPA utilizes a flexible and modular architecture where multiple oligonucleotide/oligosaccharide-binding (OB) domains are connected by flexible linkers^1,3–6^. These OB-domains harbor well defined DNA binding and protein-interaction activities^2,7^. RPA is a heterotrimer composed of RPA70 (RFA1), RPA32 (RFA2), and RPA14 (RFA3) subunits that are structurally configured as six oligonucleotide/oligosaccharide-binding (OB) domains (**Figure 1a**). The OB-domains are functionally classified as DNA binding domains (DBDs) or protein-interaction domains (PIDs). DBDs-A, B, and C reside in RPA70 and DBD-D in RPA32, respectively. PID^70N^ is situated at the N-terminus of RPA70 and tethered to DBD-A through a long-disordered F-A linker. Similarly, PID^32C^ is situated at the C-terminus of RPA32 and also connected to DBD-D by a disordered D-wh linker. PID^32C^ is also commonly known as the winged-helix (wh) motif in RPA. RPA binds to ssDNA with high affinity (K_D_ < 1 nM) and all four DBDs contribute to this property^1,2^. DBDs-A and B are more dynamic compared to DBDs-C and D^7–10^. Domains C, D, and E interact and form the trimerization core (Tri-C) that holds the RPA complex together^3^. The versatile functions of RPA are accomplished through configurational plasticity of the individual domains where differential arrangements of the domains and/or the linkers result in structures that are tailored to serve context-specific functions^2,5,11,12^. We recently showed that such configurational changes are further regulated through phosphorylation^9,10^.

**Figure 1.**
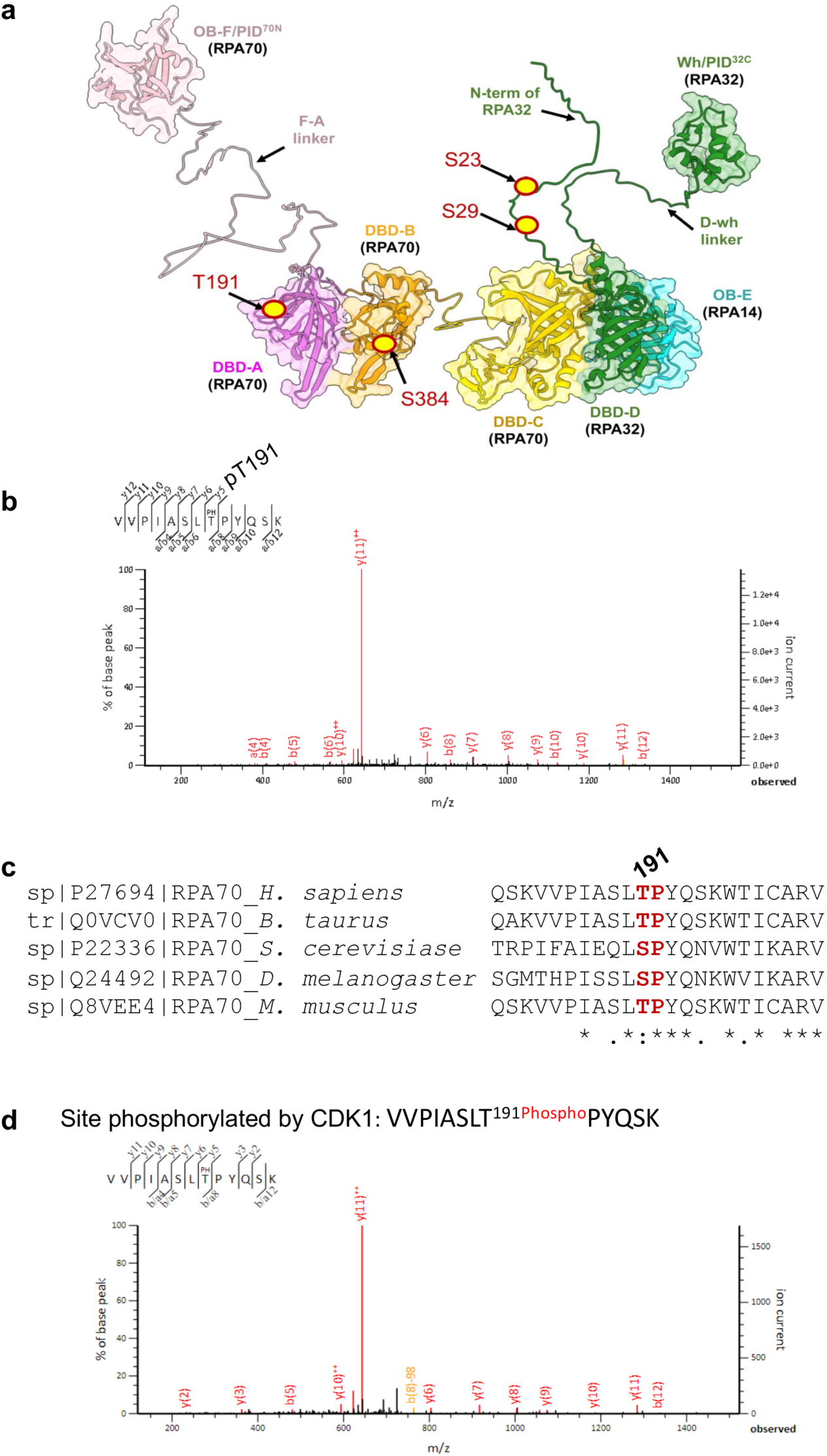
Phosphorylation of the RPA70-T191 site by Cdk1. **a.** A structural model of human RPA was generated using AlphaFold and aligned to the trimerization core observed in the crystal structure (PDB:1L1O). The RPA70, RPA32, and RPA14 subunits form a heterotrimer and harbor multiple oligosaccharide/oligonucleotide binding (OB) domains. A, B, C, and D, are DNA binding domains (DBDs). OB-F and the wh-domain are two protein-interaction domains. The cell cycle-specific sites of phosphorylation on RPA32 and RPA70 are denoted in red/yellow. **b.** Mass spectrum of the RPA70 tryptic peptide showing phosphorylation at T191 residue. MS analysis was performed using endogenous RPA70 immunoprecipitated from HCT116 cells arrested in mitosis using 50 ng/mL nocodazole for 18 hrs. Sequence alignment of RPA70 (RFA1) protein reveals a conserved SP/TP motif. (*) indicate positions which have a single, fully conserved residue, (:) indicates conservation between groups of strongly similar properties, and (.) indicates conservation between groups of weakly similar properties. **d.** Mass spectrum of the RPA70 tryptic peptide showing phosphorylation at the T191 residue. Reactions were extracted from *in vitro* kinase assay of recombinant human RPA incubated with Cdk1/Cyclin B complex followed by MS-MS analysis.

Most of the phosphosites on RPA map to a ∼40 aa disordered region in the disordered N-terminus of RPA32^13,14^. RPA32 is phosphorylated during an unperturbed cell cycle and in response to DNA damage^13,14^. During normal cell cycle progression, RPA32 is constitutively phosphorylated at the beginning of S phase, G2, and mitosis, and dephosphorylated at the end of mitosis^13–16^. Both S23 and S29 sites in the N-terminus of RPA32 have been shown to be important for cell cycle-specific phosphorylation of RPA during S and G2/M phase^13–17^. Both S23 and S29 sites harbor the canonical SP/TP motifs that are phosphorylated by the Cdk1/Cyclin B complex^13–20^. Cdk1 phosphorylates non-chromatin bound soluble RPA and the efficiency of RPA phosphorylation is independent of RPA-ssDNA interactions^17,19^. Expression of a phosphodead S23A/S29A RPA mutant hinders normal cell cycle progression by inducing a marked delay in G2^21^. However, the mechanisms that lead to this delay remain unexplored.

In addition to cell cycle-specific phosphorylation, RPA32 is also hyperphosphorylated in response to DNA damage at S4, S8, S11/12/13, T21, and S33 sites^13,14,21–33^. Phosphorylation at S33 is mediated by ATR, whereas the other sites are primarily phosphorylated by DNA-PK and/or ATM kinase^13,14,21–32^. ssDNA binding is essential for the hyperphosphorylation of RPA32 in response to DNA damage^13,26^. The critical function of hyperphosphorylation of RPA32 is to modulate DNA repair through homologous recombination (HR) and to mediate context-specific checkpoint response^13,14,21,29,31,34–37^. Phosphorylation at the S23 and S29 sites are also critical for priming the DNA damage-induced hyperphosphorylation of RPA32^13,14,21,22^. Evidence for communication between the priming and hyperphosphorylation sites on RPA32 arise from studies that show a marked decrease in hyperphosphorylation when S23 and S29 are substituted with alanine, or when treated with roscovitine, a CDK inhibitor^21,22^. These studies show that the priming phosphorylation at the S23 and S29 sites during normal cell cycle progression is critical for hyperphosphorylation of RPA32. However, it remains unresolved as to how priming at the S23 and S29 phosphosites dictate the sequence of hyperphosphorylation events and recruitment of the associated kinases.

In addition, while phospho-modified residues have been identified in both the RPA70 and RPA32, canonical functional and regulatory features have almost always been viewed through the lens of phosphorylation of RPA32. The importance of considering phosphosites in RPA70 for functional interrogation was highlighted in our recent study where we uncovered a Ser-384 site as a site for phosphorylation by Aurora B kinase^9^. RPA70-S384 phosphorylation is required for normal mitotic progression^9^. Loss of RPA70-S384 phosphorylation resulted in a feedback inhibition of Aurora B activity that led to defects in mitotic chromosome segregation and condensation^9^. This modification also inhibited the binding of BRCA2-DSS1 suggesting that phosphorylation at RPA70-S384 contributes to the suppression of HR-mediated DNA repair in mitosis. Thus, we propose the existence of a cell cycle-specific phosphocode that requires modifications spanning both RPA70 and RPA32 subunits.

Here we reveal that this phosphocode also includes modification at a novel T191 site in RPA70. RPA70-T191 is phosphorylated by the Cdk1/Cyclin B kinase complex. A T191A phosphodead substitution results in a marked delay in cell proliferation but does not induce cell death or genomic stress. The delay is primarily observed at the G2 to M transition due to a feedback inhibition of Cdk1 activity by stabilization of the inhibitory activity of Wee1 kinase. Remarkably, loss of the Cdk1-mediated RPA70-T191 phosphorylation also markedly reduces DNA damage-induced hyperphosphorylation of RPA32. Thus, the RPA70-T191 phosphosite also acts as a priming site. Using structural mass spectrometry, we show that the priming phosphocode at T191 (RPA70), S23 & S29 (RPA32) sites increases accessibility to the disordered N-terminus of RPA32 and protein-interaction domains. These collective structural changes promote the recruitment of downstream kinases such as DNA-PK and lead to hyperphosphorylation. Thus, this study unveils the mechanism of how cell cycle-specific priming leads to hyperphosphorylation of RPA in response to DNA damage. It also reveals a novel role for RPA in modulating the normal G2-M phase progression through a unique feedback regulation of the kinases the drive these cell cycle phases.

## RESULTS

### T191 in RPA70 is phosphorylated by Cdk1

To better understand RPA70 regulation in mitosis, endogenous RPA70 was immunoprecipitated from mitotic cells followed by mass spectrometric (MS) analysis. MS analysis showed that the T191 site in RPA70 is also phosphorylated in mitosis (**Figure 1b**). Phosphorylation of T191 has also been captured by other global phosphoproteomic studies including those focused on mitotic sites of phosphorylation^38–41^. These findings suggest that T191 is a novel site of phosphorylation in RPA70. T191 lies in the DBD-A domain of RPA70 (**Figure 1a**) and is highly conserved (**Figure 1c**). The T191 site also harbors the canonical SP/TP motif recognized by the Cdk1/Cyclin B kinase complex that is active during the G2 to M phase of cell cycle. A previous study had shown that RPA70 is also phosphorylated *in vitro* by Cdk1^19^. However, the potential sites of phosphorylation were not uncovered. We, therefore, performed an *in vitro* kinase assay using Cdk1 followed by MS analysis that identified T191 as a site phosphorylated by Cdk1/Cyclin B on RPA70 (**Figures 1d**). However, we did not observe T191 phosphorylation by Cdk2/Cyclin A complex by MS analysis (**Figure S1a**).

### Loss of RPA70-T191 phosphorylation delays G2 to M transition

To determine the physiological relevance of T191 phosphorylation of RPA70, we generated a homozygous knock-in T191A phosphodead mutant using CRISPR/Cas9-gene editing of HCT116 cells. Comparison of the isogenic wildtype (WT) lines with the RPA70-T191A mutant showed that the mutant grows significantly slower than WT as shown by MTS assay (**Figure 2a**). However, we did not observe any induction of cell death as determined by cleaved caspase 3 activity and morphological analysis (**Figure 2b**). The mutant cells also did not exhibit basal genomic stress as determined by an absence of change in basal p53 levels and Chk1-S317 phosphorylation (**Figure 2c**). Given that Cdk1/Cyclin B activity is integral to the G2 to M transition, we also assayed for changes in mitotic population of the T191A mutant. Remarkably, the percentage of mitotic cells in the T191A mutant are significantly lower than the WT cells as shown by reduced phospho-S10- Histone H3 staining (**Figures 2d** and **e**), which suggests a potential delay in the G2 to M phase. Analysis of overall cell cycle profiles using flow cytometry also shows a mild increase in the G2/M population, though this approach does not distinguish G2 from M-phase (**Figure S1b**).

**Figure 2.**
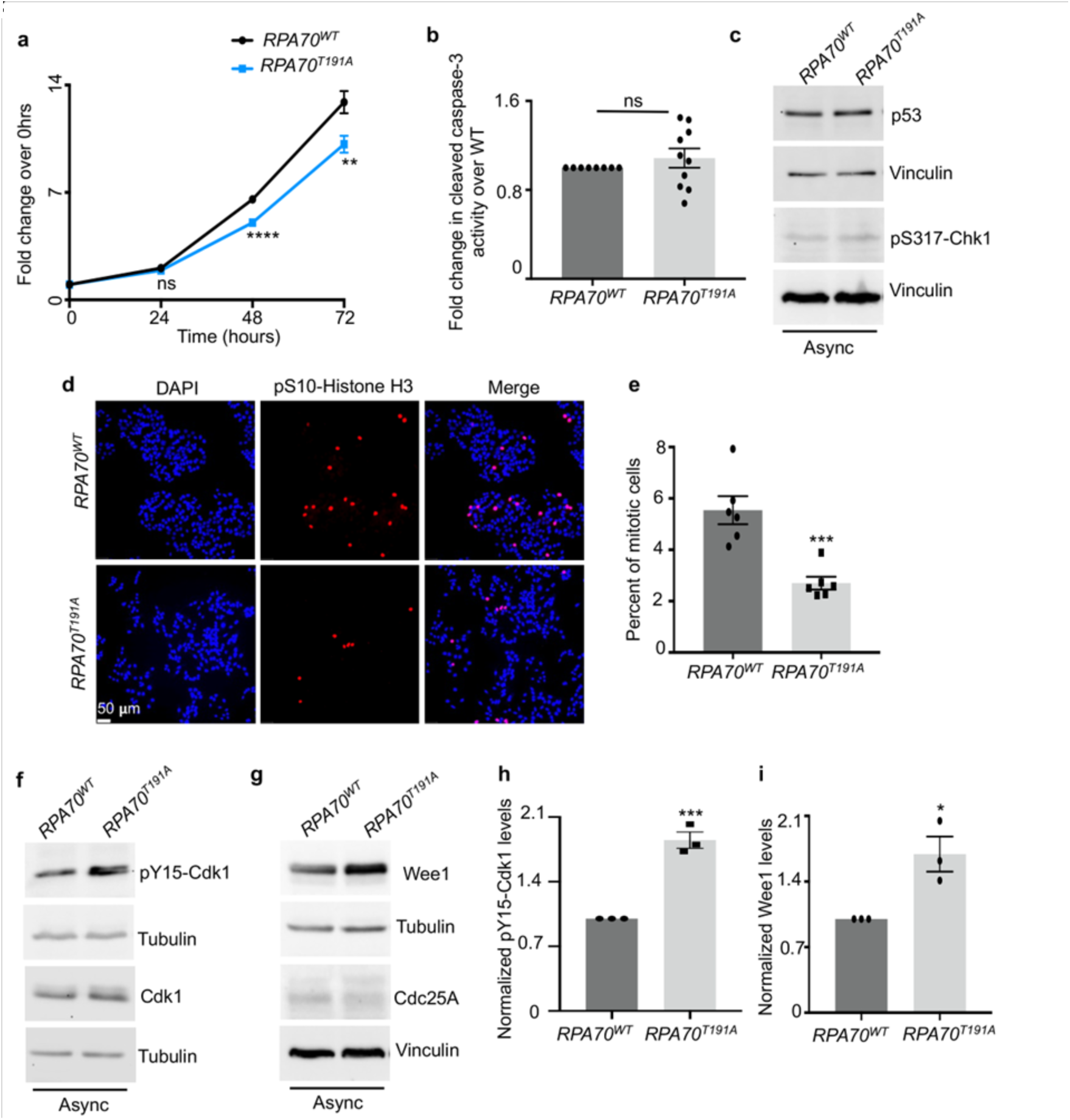
**a. Decrease in mitotic population of phosphodead mutant is due to inhibition of Cdk1 activity and stabilization of Wee1**. MTS assay shows decreased growth rate of the mutant. Cells were assayed at 0, 24, 48, and 72 hrs of growth. Values corrected for background absorbance were normalized to 0 hrs of growth. Error bars denote SEM. The mean of three independent experiments are plotted. Triplicate wells were assayed per time point for each experiment. Statistical significance was determined using an unpaired two-tailed *t*-test: *p* = 0.08 at 24 hrs, (ns = not significant), \*\*\*\**p* = 0.00006 at 48 hrs, ** *p* = 0.008 at 72 hrs. **b.** Caspase 3/7 activity was determined by the Caspase3/7 glo assay. Bar graph depicts data corrected for background absorbance and plotted as a fold change over WT-control. Error = SEM. The mean of three independent experiments are plotted. 3 to 4 wells were assayed per experiment. Statistical significance was determined using an unpaired two-tailed *t*-test: *p* = 0.38 (ns). **c.** Western blot represents p53 and phospho-S317-Chk1 levels in asynchronous (Async) WT and mutant HCT116 cells. Vinculin was probed as loading control. Data represents three independent experiments. **d**. Representative immunofluorescent images of asynchronous cells stained with DAPI (nuclear marker) and anti-phospho-S10-Histone H3 antibody (mitosis). **e.** Percent of mitotic cells stained red in the images shown in d. were quantitated using six image panels with an average of 350 cells per panel from three independent experiments. Error bars denote SEM and significance determined by an unpaired two-tailed *t*-test: *** *p* = 0.0008. **f and g.** Western blot analysis of asynchronous cells probed with indicated antibodies. Tubulin and Vinculin were probed as loading controls. **h and i.** Plots represent Cdk1–Y15 phosphorylation levels and Wee1 levels normalized to the loading control. Data are representative of three experiments. Error bars=SEM. Statistical significance determined by unpaired two-tailed *t*-test: \*\*\**p* = 0.0007 (h) and \**p* = 0.0208 (i).

### RPA70-T191 phosphorylation maintains Cdk1/Wee1 activity

To better understand the delay in G2 to M transition, we assessed the changes in Cdk1 activity. G2 to M transition is initiated by formation of the Cdk1/Cyclin B complex and by activating phosphorylation at the T161 site on Cdk1^42–47^. The key regulatory step in the activation of Cdk1 is the removal of inhibitory phosphorylation at the T14/Y15 sites. Wee1 is the major tyrosine kinase that phosphorylates the Y15 site and inhibits Cdk1 activity^43,48–53^. In late G2 phase, the Cdc25 family of phosphatases remove the inhibitory T14/Y15 phosphorylation to activate Cdk1^44,45,49,51,52,54–60^. Once Cdk1 is fully active, it phosphorylates Wee1 and targets it for degradation^61–63^. Cdk1 also phosphorylates and maintains the stability of Cdc25A^44^. This creates a positive feedback loop wherein Cdk1 activity is maintained through degradation of Wee1 and stabilization of Cdc25A that permits entry into mitotic phase.

Given the decrease in mitotic cells, we probed for changes in Y15 phosphorylation of Cdk1 in asynchronous cells (**Figure 2f**). We found that Y15 phosphorylation of Cdk1was significantly increased by more than 2-fold in the mutant (**Figures 2f** and **h**) and this increase was not attributed to changes in total Cdk1 levels (**Figure 2f**). The total Wee1 levels were also markedly increased in the mutant cells indicating that stabilization of Wee1 led to an increase in the inhibitory Y15 phosphorylation of Cdk1 (**Figures 2g, h** and **i**). We also probed for changes in levels of Cdc25A, which is a labile protein with a very short half-life that becomes stabilized in late G2 to M phase^64,65^. The basal levels of Cdc25A in asynchronous cells were quite low but were not altered in the mutant relative to WT (**Figure 2g**). This suggests that the decrease in mitotic population is due to a delay in G2 to M transition mediated by stabilization of Wee1 that in turn inhibits Cdk1 activity.

To further assess the G2 to M delay, we synchronized the cells using double-thymidine block. Cells were arrested at the G1/S transition in both WT and the mutant (**Figure S2a**). Once the cells were released, there were no substantial delays in progression through S-phase and early to mid G2 phase (**Figure S2b**). Since Cdk1/Wee1 axis primarily regulates the G2 to M transition, we probed for changes in Wee1 activity in late G2 phase and as they entered mitosis. Wee1 levels were found to be markedly induced at the G2 to M transition in the mutant cells (**Figure 3a**). We found that the inhibitory Y15 phosphorylation of Cdk1 was markedly induced in late G2 but not in early M phase (**Figure 3a**). The RPA70-T191A mutant cells were able to enter mitosis, however, the RPA70-T191A cells showed delayed mitotic entry as determined by low levels of S10 phosphorylation of Histone H3 in the mutant cells relative to WT at the same time point (**Figure 3a**). This further indicated a delay in the G2 to M transition in the mutant. However, we did not observe changes in the total levels of Cyclin B, Cdk1, Cdc25A, or RPA70 (**Figure 3b**), further suggesting that the loss of RPA70-T191 phosphorylation primarily affects Wee1 levels and thereby Cdk1-Y15 phosphorylation.

**Figure 3.**
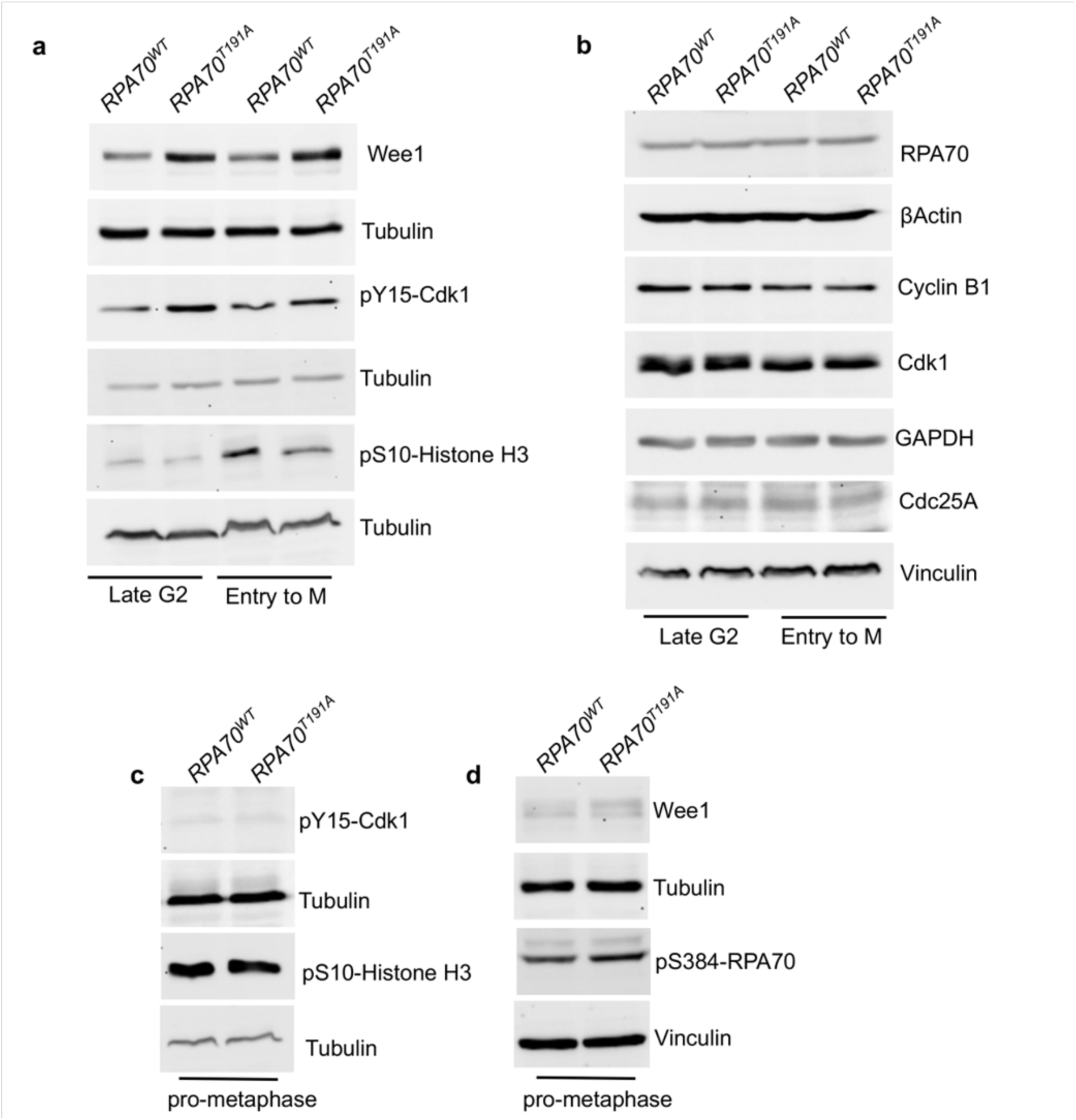
a. and b. Increased levels of Wee1 and Y15-phosphorylation of Cdk1 in the RPA70- T191A mutant delay entry into mitosis. HCT116 cells, synchronized by double thymidine arrest, were released, and cells collected at 7 hrs and 8.5 hrs post-release representing late G2 phase and entry into Mitotic (M) phase, respectively. Western blots were probed with the indicated antibodies. Entry into M was validated with increased phospho-S10 Histone H3 levels. Blots are representative of three independent experiments. Vinculin, Tubulin and βActin were probed as loading controls. **c and d.** Western blot analysis of mitotic cells arrested in pro-metaphase using nocodazole (75ng/mL) for 18 hrs and collected by shake-off method. Blots were probed with the indicated antibodies. Tubulin and Vinculin were probed as loading controls. Blots are representative of three independent experiments.

### Loss of RPA-T191 phosphorylation does not affect progression through mitosis

To determine if the mutant cells exhibited defects in Cdk1 activity as they progressed through mitosis, cells were synchronized in pro-metaphase using nocodazole. Progression through mitosis was not hindered as shown by similar levels of S10 phosphorylation of Histone H3 in both WT and mutant cells (**Figure 3c**). Also, the expected loss of Y15-Cdk1 phosphorylation and degradation of Wee1 kinase in mitosis were observed to similar extent in both WT and mutant cells (**Figure 3c** and **d**). We previously showed that decrease in RPA70-S384 phosphorylation markedly affected mitotic progression by inhibiting Aurora B kinase activity that led to defects in chromosome segregation and condensation^9^. However, we did not observe any such decrease in the mitosis-specific RPA70-S384 phosphorylation (**Figure 3d**). Furthermore, we did not observe any defects in chromosome segregation or condensation (**Figures S3a** and **b**). No changes in spindle assembly were observed (**Figures S3a** and **b**). These results suggest that although loss of RPA70-T191 phosphorylation delays G2 to M transition, once these cells enter mitosis, they are able to complete mitosis without impediments. Thus, these findings have uncovered a unique feedback regulation of Cdk1 activity and Wee1 levels by RPA through Cdk1-dependent phosphorylation of RPA70 that primarily regulates the G2 to M transition.

### T191 phosphorylation at RPA70 is important for priming hyperphosphorylation of RPA32 in response to DNA damage

A hallmark of the Cdk1-regulated sites on RPA32 (S23 and S29) is that it primes the hyperphosphorylation of RPA32 at the S4/8, S33, S11/12/13, and T21 sites in response to DNA damage (**Figure 4a**). We therefore wanted to determine if the Cdk1-regulated T191 phosphosite on RPA70 is also critical for hyperphosphorylation of RPA32 in response to DNA damage. To induce DNA damage, cells were briefly incubated with SN-38, a topoisomerase I inhibitor and an active metabolite of irinotecan. Intriguingly, the RPA70-T191A mutant cells were markedly sensitive cells to genomic stress and displayed higher levels of apoptosis in response to SN-38 treatment (**Figure 4b**). To better understand this enhanced sensitivity to genomic stress, we also probed for changes in hyperphosphorylation of RPA32. Remarkably, loss of T191 phosphorylation markedly affected the hyperphosphorylation of RPA32 as indicated by a decrease in the S33 phosphorylation and S4/S8 phosphorylation relative to WT (**Figure 4c**).

**Figure 4.**
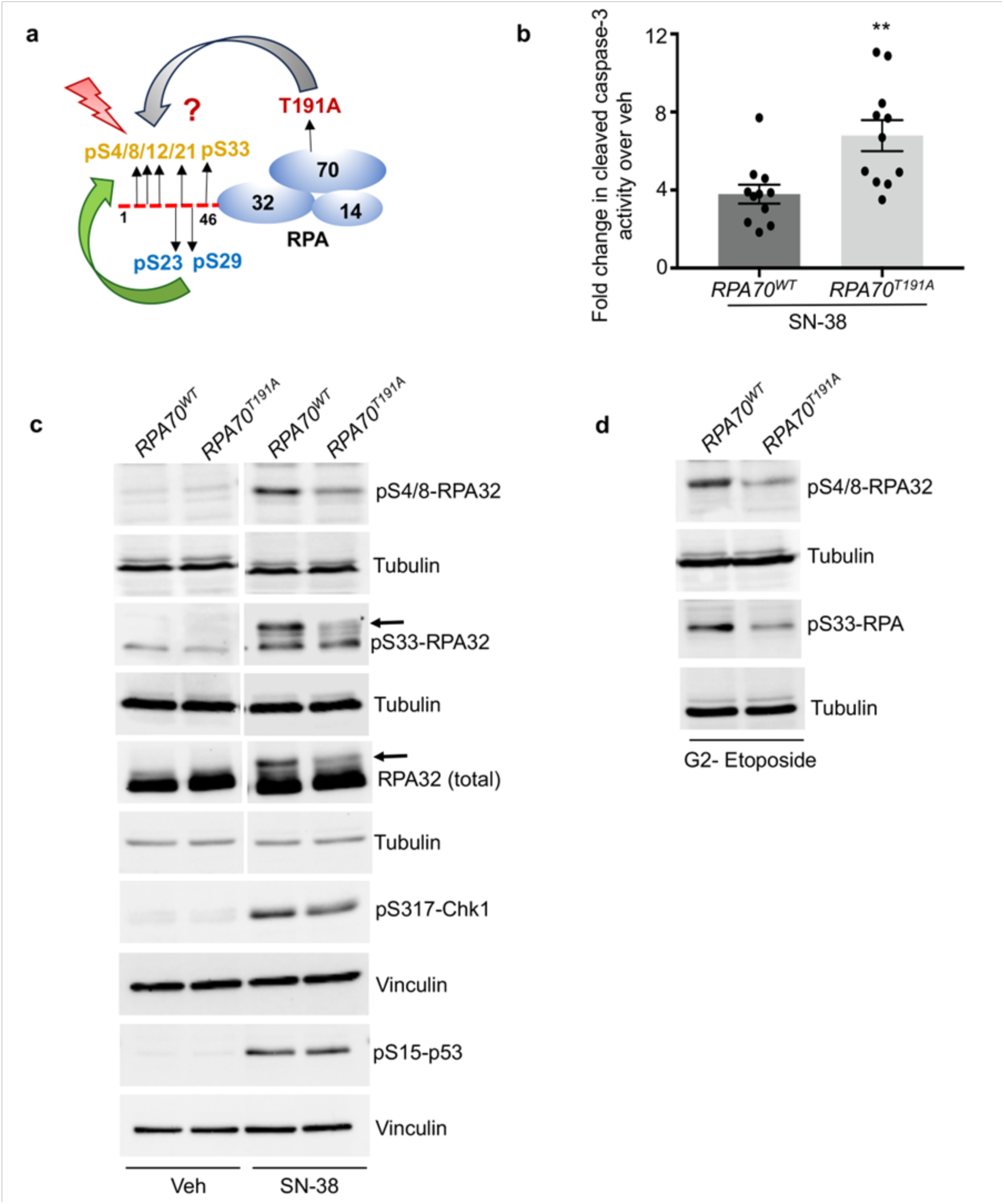
Loss of T191 phosphorylation in RPA70 enhances sensitivity to DNA damage by limiting hyperphosphorylation of RPA32. **a.** Cartoon depicts hyperphosphorylation of the N-terminus of RPA32 in response to DNA damage. It is unclear as to how disruption of T191-RPA70 phosphorylation affects hyperphosphorylation of RPA32. Cdk1-primed sites (phospho-S23 and S29) on RPA32 are indicated in blue. **b.** Caspase 3/7 activity in cells treated with 100 ng/mL SN- 38 for 21hrs was determined by the Caspase3/7 glo assay. Bar graph depicts data corrected for background absorbance and plotted as fold change over vehicle (veh) control (0.1% DMSO). Error = SEM. Mean of three independent experiments are plotted. 3 to 4 wells were assayed per experiment. Statistical significance was determined using an unpaired two-tailed *t*-test: \*\*\**p* = 0.0042. **c.** Western blot analysis of cells treated briefly with 20 ng/mL SN-38 for 90 min or vehicle (0.1% DMSO). Blots were probed with the indicated antibodies with Tubulin and Vinculin as loading controls. Blots are representative of three independent experiments. **d.** Cells synchronized with double thymidine block were released and at 7 hrs post-release, cells in G2 were treated with 100 μM etoposide for 45 min. Blots were probed with the indicated antibodies with Tubulin as loading control. Blots are representative of three independent experiments.

Hyperphosphorylation of RPA32 has been shown to affect cell cycle checkpoint responses, but it can vary depending on the cellular context^13,14^. However, we did not observe marked differences in Chk1-S317 phosphorylation or p53-S15 phosphorylation suggesting that Chk1 and p53 checkpoint responses were intact in the mutant cells (**Figure 4c**). Similarly, cells arrested in late G2, followed by a brief incubation with etoposide (a topoisomerase II inhibitor), also exhibited a marked decrease in the DNA damage-induced hyperphosphorylation of RPA32 (**Figure 4d**). Both S4/S8 and S33 phosphorylation were markedly reduced in the mutant cells in G2 phase in response to DNA damage (**Figure 4d**). The S33 phosphorylation is usually observed both in the basal form of RPA32 and in the hyperphosphorylated form in response to DNA damage^13^. However, we did not observe a marked increase in the slower migrating species of RPA32. Instead changes in S33 phosphorylation were observed only in the basal phosphorylated form in the G2 phase (**Figure 4d**). Such variations in the extent of RPA hyperphosphorylation in response to DNA damage based on cell cycle phase and DNA damaging agent, have been observed before^13^. Thus, our results have uncovered the Cdk1-regulated T191 site on RPA70 as an additional critical regulator of hyperphosphorylation in synergy with the other Cdk1-priming sites (S23 and S29) on RPA32. It is also highly likely that the effect on hyperphosphorylation of RPA32 introduced by the T191A mutation is due to inhibition of Cdk1 activity that would in turn diminish S23 and S29-priming phosphorylation. These results also show for the first time that a phosphorylation-dependent regulation of the large RPA70 subunit can in turn regulate the RPA32 subunit. Thus, these results emphasize an extensive inter-subunit, phosphorylation-driven, regulatory network within RPA.

### A phosphomimetic T191D substitution does not alter the secondary structure or ssDNA binding properties of RPA

To test whether phosphorylation at the T191 position influences the DNA binding properties of RPA, we recombinantly purified RPA carrying a phosphomimetic T191D substitution in the RPA70 subunit (RPA^T191D^; **Figure 5a**). As a control, we also purified RPA with a T191A substitution and investigated the ssDNA and oligomerization properties. In circular dichroism (CD) spectroscopy analysis, RPA, RPA^T191D^, and RPA^T191A^ show similar secondary structure profiles with minimal differences suggesting that the core structures of the OB-domains are not altered (**Figure 5b**). We next tested the ssDNA binding activity of all three proteins by following the change in anisotropy of fluorescein-labeled ssDNA as a function of RPA concentration. The site-size of ssDNA binding to RPA is ∼18-24 nt^10,66^ and thus we tested DNA binding using two short poly-dT oligonucleotides of varying lengths. RPA stoichiometrically bound to both (dT)_20_ or (dT)_35_ substrates with high affinity and we observed no differences between RPA, RPA^T191D^, or RPA^T191A^ proteins (**Figures 5c** and **d**). We further probed the kinetics of RPA-ssDNA interactions using a stopped flow assay where the changes in intrinsic Trp fluorescence of RPA were monitored. Several Trp residues in RPA base-stack with the ssDNA resulting in a quenching of fluorescence signal. Performing these experiments as a function of ssDNA concentration allowed us to compare the kinetics of ssDNA binding. RPA, RPA^T191D^, and RPA^T191A^ all bound to ssDNA with rapid and comparable k_on_ values (**Figures 5e-h**). These data suggest that there are no significant differences in the ssDNA binding properties upon phosphorylation at the T191 position.

**Figure 5.**
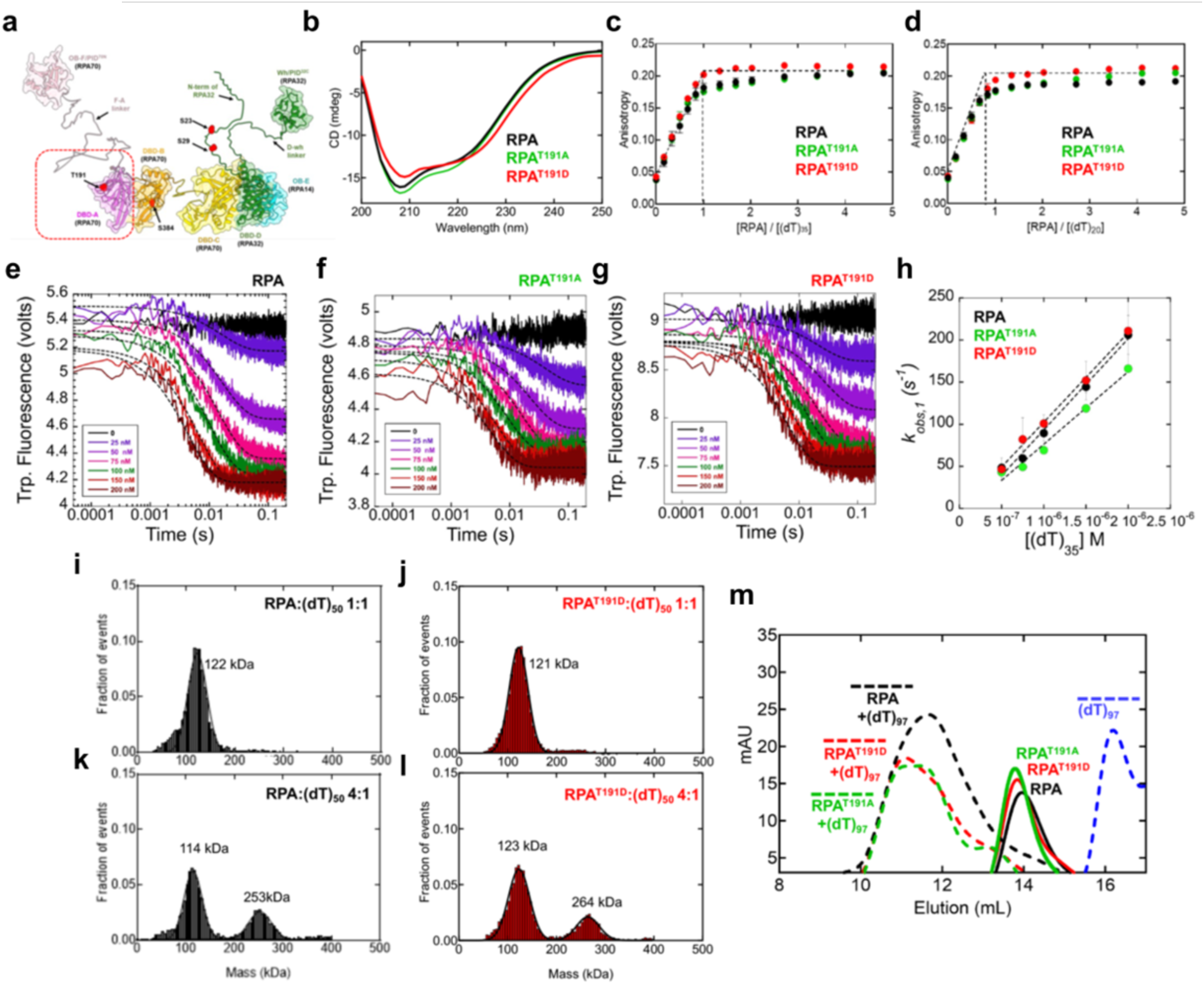
Phosphorylation of RPA-T191 does not alter the secondary structure or ssDNA binding properties. **a.** Structural model of the domains of RPA modeled using AlphaFold. The positions of phosphorylation (T191 and S384 in RPA70 and S23 and S29 in RPA32 are depicted. **b.** Circular dichroism (CD) spectra of RPA, RPA-T191A, and T191D does not show significant differences in the overall secondary structures. ssDNA binding activity of RPA, RPA-T191A, and RPA-T191D were measured using fluorescence anisotropy and fluorescein-labeled **c.** (dT)_35_ or **d.** (dT)_20_ oligonucleotides. ssDNA binding of all three RPA proteins are stoichiometric due to the high-affinity ssDNA binding activity of RPA. Stopped flow analysis of RPA-ssDNA interactions were performed by following the change in intrinsic Trp fluorescence changes as a function of increasing (dT)_35_ concentrations. Data for **e.** RPA, **f.** RPA-T191A, and **g.** RPA-T191D are shown. Data were fit to single exponentials and **h.** the k_obs_ values plotted again ssDNA concentration yields similar k_ON_ values for ssDNA binding for all three RPA proteins (RPA = 1.08±0.2×10^8^M^-1^s^-1^, RPA- T191A = 0.86±0.3×10^8^M^-1^s^-1^, RPA-T191D = 1.06±0.2×10^8^M^-1^s^-1^). **i-l.** Mass photometry analysis of RPA or RPA-T191D complexes on (dT)_50_ ssDNA were performed at either 1:1 or 4:1 molar ratios of RPA:DNA. Predominantly single RPA-bound ssDNA complexes are observed at 1:1 ratios and both 1:1 and 2:1 RPA-bound complexes are observed. However, there are no quantifiable differences in the ssDNA binding properties between RPA and RPA-T191D. **m.** Size exclusion chromatography analysis of RPA, RPA-T191A, and RPA-T191D in the absence of presence of longer (dT)_97_ ssDNA substrates show similar profiles for RPA interactions. In these experiments, a three-fold excess of RPA was used to drive the assembly of multiple RPA molecules on the DNA. For all these experiments, representative data from a minimum of three independent experiments are shown. Error bars, where appropriate, are shown and represent St. Dev. values from n ≥ 3 independent experiments.

Since the ssDNA binding properties were not influenced by the phosphomimetic modification at T191, we next tested whether assembly of multiple RPA molecules were modulated. On long ssDNA substrates, unmodified RPA does not interact with cooperativity. However, in our previous analysis of a phospho-modification at Ser-384 in RPA70, a higher density of RPA^S384D^ occupied the ssDNA compared to unmodified RPA^9^. Thus, we next tested the binding of multiple RPA or RPA^T191D^ molecules to a (dT)_50_ substrate under stoichiometric (1:1 ssDNA:RPA) or sub stoichiometric (1:4 ssDNA:RPA) molar ratios using mass photometry. In experiments with 1:1 RPA:DNA, one molecule of RPA or RPA^T191D^ engage the DNA (**Figures 5i** and **j**). Under conditions of excess RPA, both one and two RPA-bound complexes are captured, but there are no observed differences between RPA versus RPA^T191D^ (**Figures 5k** and **l**). Analysis of RPA binding to longer (dT)_97_ ssDNA substrates were performed using size exclusion chromatography under 3:1 ratios of RPA:DNA. Even under these conditions, no differences in complex formation are observed between RPA and RPA^T191D^ (**Figure 5m**). RPA^T191A^ was also assessed as a control and showed ssDNA binding behavior similar to RPA. Thus, the *in vivo* functional differences observed are likely not attributable to changes in ssDNA binding properties.

RPA is phosphorylated by Aurora B kinase during mitosis at Ser-384 in RPA70 (**Figure 5a**)^9^. This modification resulted in configurational rearrangement of the individual domains and led to formation of higher density RPA-ssDNA complexes. Thus, we next tested whether RPA^T191D^ had similar effects. We first assessed the intrinsic Trp fluorescence at equal protein concentrations (100 nM each) and find that the fluorescence of RPA^T191D^ is about half that of RPA (**Figure 6a**). Trp fluorescence is sensitive to the polarity of the environment and under polar conditions a reduction in fluorescence is indicative of exposure of the Trp residues to solvent^67^. This fluorescence reduction occurs due to dissipation of the excited-state energy to the solvent. Thus, an unfolding or opening/remodeling of RPA^T191D^ is reflected by the reduction in Trp fluorescence.

**Figure 6.**
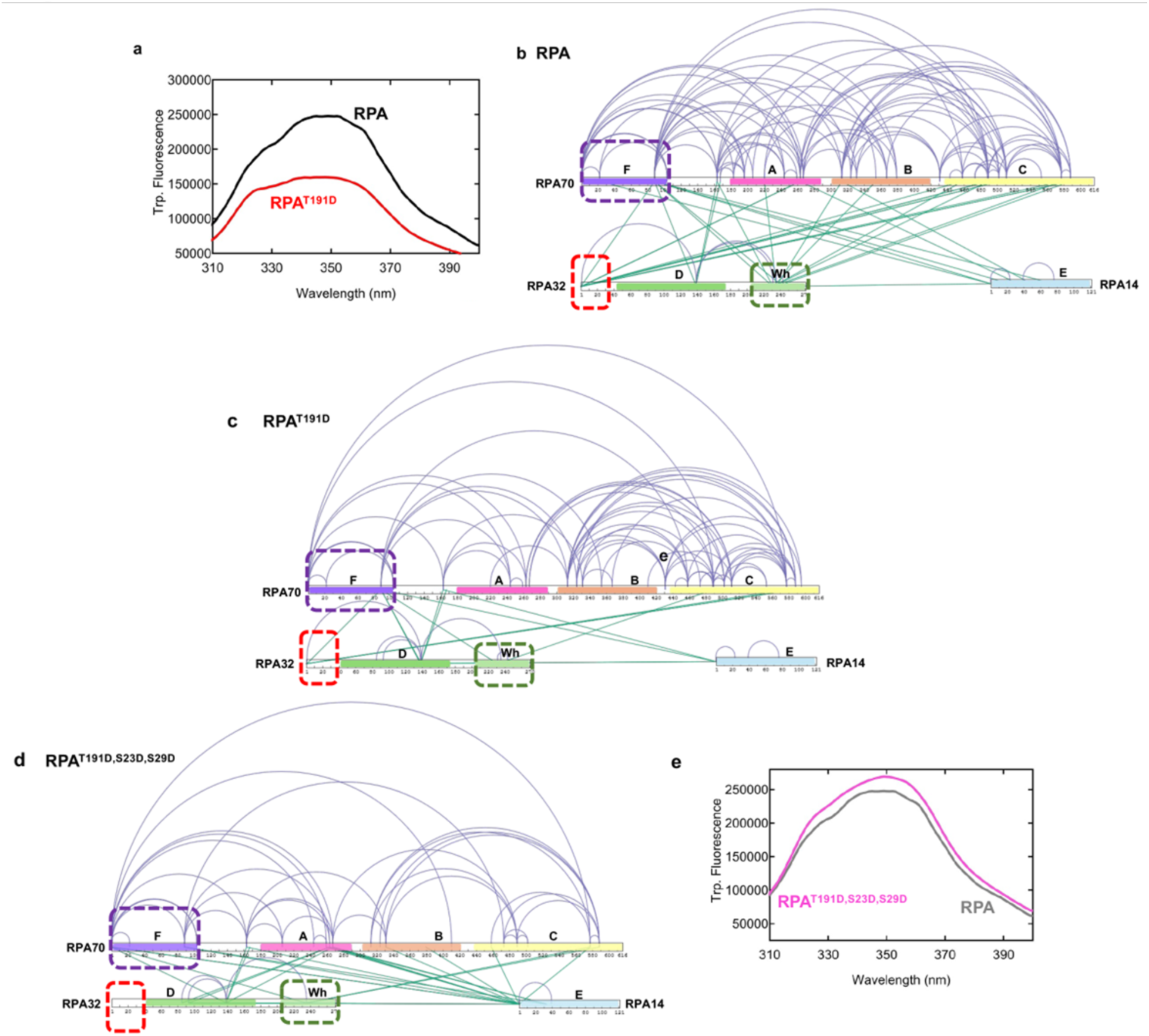
Differential configuration changes in RPA are introduced through the phosphocode. **a.** Intrinsic Trp fluorescence scan of RPA versus RPA-T191D shows reduced signal for the phosphomimetic mutant. **b.** Crosslinking mass spectrometry analysis of RPA with BS3. Crosslinks (XLs) within each subunit and between the three subunits are observed. The F-E OB domains and the wh domain are denoted. The disordered N-terminus of RPA32 and two protein interaction domains (F and wh) are marked by the dashed squares. **c.** XL-MS analysis of RPA- T191D shows lesser overall XLs compared to RPA in panel b. In particular, XLs originating from the two protein interaction domains (F and wh) and the N-terminus of RPA32 are all reduced. XLs in RPA14 are also reduced. Data suggest an overall configurational opening of RPA-T191D. **c.** XL-MS analysis of the RPA-T191D,S23D,S29D triple phosphomimetic mutant shows pattern of crosslinking that are different compared to RPA or RPA-T191D. No XLs are captured in the N-terminus of RPA32 suggesting that this region is now fully accessible (dashed square). **e.** Intrinsic Trp fluorescence scan of RPA-T191D,S23D,S29D show differences that are marginally higher than RPA. This observation is different that the intrinsic Trp profile for RPA-T191D (panel a). These data show defined configurational states driven by the phosphocode. For all these experiments, representative data from a minimum of three independent experiments are shown.

### Structural mass spectrometry reveals rearrangements of the protein interaction domains of RPA upon phosphorylation at T191

To obtain more structural information, with respect to remodeling of the domains of RPA versus RPA^T191D^, we performed crosslinking mass spectrometry (XL-MS) after treating the proteins with bis(sulfosuccinimidyl)suberate (BS3). BS3 crosslinks primary amines that are within ∼11-20 Å^68^ and thus can provide a coarse-grain model of the relative positions of the various domains in RPA^69^. Cdk1 phosphorylates RPA that is not bound to DNA and the efficiency of phosphorylation is not stimulated by ssDNA^17,19^. Thus, the following experiments were performed in the absence of ssDNA. For RPA and RPA^T191D^ samples, an extensive number of crosslinks (XLs) are captured between the F, A, B, and C domains within RPA70 (**Figures 6b** and **d**). XLs between RPA70, RPA32, and RPA14 are also captured. For unmodified RPA, a structural organization can be envisioned where OB domains-A, B, C, D and E are in a ‘C-shaped’ configuration with OB-F and the wh-domains situated on top. Such a model is predicted by AlphaFold-3 (**Figure S4**) and supported by the structural and biochemical studies of the RPA^6,10,70^. However, the first difference is that the number of XLs within RPA70 and between the three subunits are fewer in the RPA^T191D^ sample (**Figure 6c**).

For clarity, we here focus on XLs that originate from OB-F (PID^70N^), Wh (PID^32C^), and the N-terminus of RPA32. In the RPA^T191D^ dataset, XLs from OB-F to OB-A, B, and C are lost. XLs from the RPA32 N-terminus to OB-A, and XLs to the 410-500 region are lost. Finally, XLs from the Wh-domain to the F-A linker, OB-A, OB-B, and almost all (except one) to OB-C are lost. These XL-MS analyses support the Trp fluorescence findings and suggest that the protein-interaction domains are released from a compacted configuration of RPA. These data suggest that the protein-interaction domains and the N-terminus of RPA32 become more accessible upon phosphorylation at T191 by Cdk1.

### A Cdk1-specific phosphocode drives further configurational rearrangements of RPA

In addition to the new RPA70-T191 site identified in this study, Cdk1 is known to phosphorylate S23 and S29 sites in the N-terminus of RPA32^13,14^. These modifications have been captured under both normal cell cycle and DNA damaging conditions^13,14^. Since our XL-MS analysis captured configurational changes that included both protein interaction domains and the disordered N-terminus of RPA32 (**Figure 6**), we wondered if Cdk1 introduced a set of phosphocode-driven configurational changes that might be additive in nature. Thus, we probed the XLs in a triple phosphomimetic RPA^T191D,S23D,S29D^ protein. Remarkably, in RPA^T191D,S23D,S29D^, all the XLs in the N-terminus of RPA32 are lost, suggesting that this region is now fully accessible (**Figure 6d**). This region in RPA is hyperphosphorylated by a slew of kinases including ATM, DNA-PK, and ATR^13,14^. In addition, new XLs between DBD-D and DBD-A are captured along with a host of new XLs between RPA14 (OB-E) and OB-F, A and B domains (**Figure 6d**). The OB-F and the Wh domains remain accessible, however, the F-domain is repositioned closer to RPA14 compared to RPA^T191D^. Corresponding to this repositioning of OB-F, the intrinsic Trp fluorescence is restored to a level slightly higher than RPA (**Figure 6e**). These changes are specific to RPA^T191D,S23D,S29D^, as the intrinsic Trp fluorescence of the RPA^S23D,S29D^ protein resemble RPA^T191D^ (**Figure S5a**). Thus, individual phosphosites remodel RPA that are structurally distinct from RPA^T191D,S23D,S29D^. However, the overall structural consequence of remodeling upon phosphorylation by Cdk1 at either site, or together, elicits changes in the two protein interaction domains and the N-terminus of RPA32 to varying degrees.

### The Cdk1 phosphocode elicits subtle changes in the assembly of RPA-ssDNA complex

Since the configurational changes of RPA^T191D,S23D,S29D^ were more pronounced, we next probed whether the combined phosphomimetic substitutions altered the ssDNA binding properties. T191 is located in RPA70 whereas S23 and S29 are in the disordered N-terminus of RPA32 (**Figure 7a**). In circular dichroism analysis, RPA^S23D,S29D^ and RPA^T191D,S23D,S29D^ do not show appreciable changes in the secondary structure (**Figure 7b**). Both proteins also bind stoichiometrically to a short fluorescein-(dT)_35_ oligonucleotide with similar affinity in fluorescence anisotropy experiments (**Figure 7c**). Furthermore, stopped flow analysis of ssDNA binding kinetics do not show an appreciable difference in binding properties between the three proteins (**Figures 7d-f**). Thus, similar to our observations for RPA^T191D^, RPA^S23D,S29D^ and RPA^T191D,S23D,S29D^ are able to interact with ssDNA similar to unmodified RPA. Since the configuration of RPA^T191D,S23D,S29D^ was different compared to RPA or RPA^T191D^ (**Figure 6**), we next tested where the binding density of RPA on longer DNA substrates were altered by the phosphocode. In mass photometry analysis, both RPA^S23D,S29D^, and RPA^T191D,S23D,S29D^ formed single-RPA bound complexes on a (dT)_50_ under stoichiometric RPA:DNA conditions (**Figures 7g** and **h**). At higher ratios of RPA:DNA (4:1), both one and two RPA-bound complexes are observed. Notably, for RPA and RPA^T191D^, the fraction of the two-RPA bound species was ∼25% (**Figures 6k** and **l**). In contrast, we observed ∼50% of the two-RPA bound species for RPA^S23D,S29D^ and RPA^T191D,S23D,S29D^ (**Figures 7i** and **j**). Correspondingly, analysis of RPA binding density on a longer (dT)_97_ ssDNA substrate using size exclusion chromatography shows a larger eluting complex for RPA^T191D,S23D,S29D^ (**Figure 7k**). These results show that the triple phosphomimetic RPA enhances the number of RPA molecules assembled on longer ssDNA substrates. Since hyperphosphorylation of RPA at additional sites (S4, S8, S11/12/13/ T21, and S33) is known to be dependent on binding to ssDNA, this suggests that priming at all three sites promotes an RPA configuration on ssDNA that is likely conducive for DNA repair.

**Figure 7.**
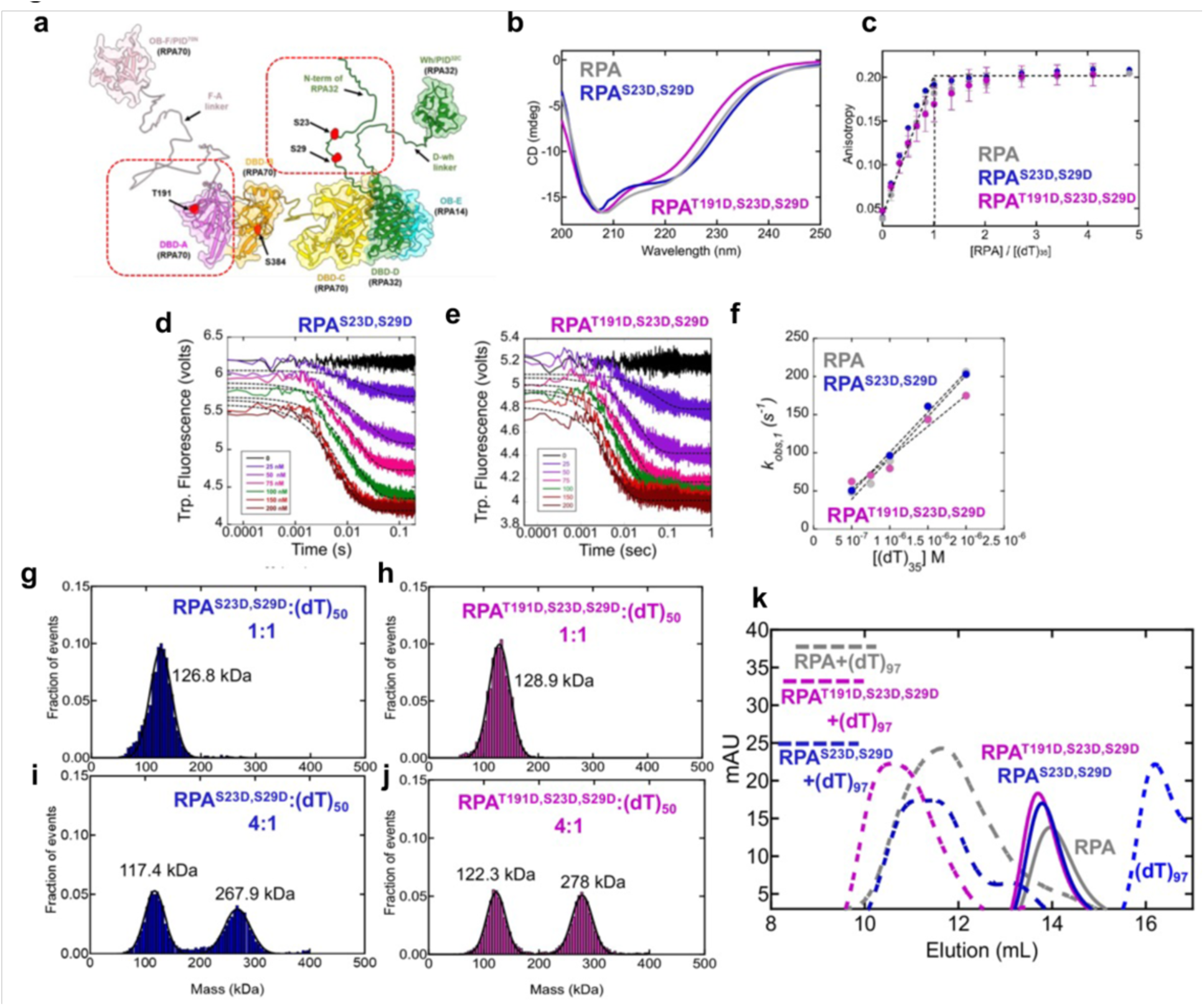
RPA-S23D,S29D does not alter the secondary structure or ssDNA binding properties, but the triple RPA-T191D,S23D,S29D phosphomimetic drives formation of higher-density RPA-ssDNA complexes. **a.** Structural model of the domains of RPA modeled using AlphaFold. The positions of phosphorylation (T191 and S384 in RPA70 and S23 and S29 in RPA32 are depicted. Red dotted boxes denote the position probed here through phosphomimetic substitutions. **b.** Circular dichroism (CD) spectra of RPA and the RPA phosphomimetic variants shows no significant differences in the overall secondary structures. **c.** ssDNA binding activity of RPA and RPA phosphomimetic variants were measured using fluorescence anisotropy and a fluorescein-labeled (dT)_35_ oligonucleotide. ssDNA binding of all three RPA proteins are stoichiometric due to the high-affinity ssDNA interactions of RPA. Stopped flow analysis of RPA-ssDNA interactions were performed by following the change in intrinsic Trp fluorescence changes as a function of increasing (dT)_35_ concentrations. Data for **d.** RPA-S23D,S29D and **e.** RPA- T191D,S23D,S29D are shown. Data were fit to single exponentials and **f.** the k_obs_ values plotted again ssDNA concentration yields similar k_ON_ values for ssDNA binding for RPA proteins (RPA = 1.08±0.2×10^8^M^-1^s^-1^, RPA-S23D,S29D = 1.06±0.2×10^8^M^-1^s^-1^, RPA-T191D,S23D,S29D = 0.85±0.1×10^8^M^-1^s^-1^). **g-j.** Mass photometry analysis of RPA or the RPA phosphomimetic variants on (dT)_50_ ssDNA were performed at either 1:1 or 4:1 molar ratios of RPA:DNA. Predominantly single RPA-bound ssDNA complexes are observed at 1:1 ratios and both 1:1 and 2:1 RPA-bound complexes are observed at higher ratios. However, for RPA-T191D,S23D,S29D there is a higher fraction of the 2:1 complex. **k.** Size exclusion chromatography analysis of RPA and the RPA phosphomimetics in the absence or presence of longer (dT)_97_ ssDNA substrates show similar profiles for RPA and RPA-S23D,S29D interactions. However, RPA-T191D,S23D,S29D for complexes that are much larger suggesting more RPA molecules bound the ssDNA for this RPA phosphomimetic variant. In these experiments, a three-fold excess of RPA or RPA- phophomimetics variants were used to drive the assembly of multiple RPA molecules on the DNA. For all these experiments, representative data from a minimum of three independent experiments are shown. Error bars, where appropriate, are shown and represent St. Dev. values from n ≥ 3 independent experiments.

### RPA remodeling by cell cycle-specific phosphocode primes hyperphosphorylation by DNA-PK

These biophysical results suggest that the cell cycle-specific phosphocode (T191, S23, S29) enhances the accessibility of the protein interaction domains and the N-terminus of RPA32. This in turn would prime the N-terminus of RPA32 for efficient hyperphosphorylation by downstream DNA kinases such as DNA-PK in response to DNA damage. To test this model, we determined if priming at all three sites: T191 and S23/S29 enhanced the efficiency of S4/S8 phosphorylation of RPA^T191D,S23D,S29D^ by DNA-PK relative to unphosphorylated RPA. Both RPA and the mutant were subjected to *in vitro* kinase reactions with DNA-PK followed by western blot analysis. Remarkably, the extent of RPA32 phosphorylation by DNA-PK was markedly enhanced by RPA^T191D,S23D,S29D^ relative to unphosphorylated (non-primed) RPA (**Figure 8a**). This priming effect was consistent across varying concentrations of RPA. Control reactions with only the substrate or kinase do not show any non-specific bands (**Figure S5b**). These data further support our finding that the priming phosphosites enhance the accessibility of the protein interaction domains and the N-terminus of RPA32 to promote recruitment of kinases such as DNA-PK to induce hyperphosphorylation of RPA32.

**Figure 8.**
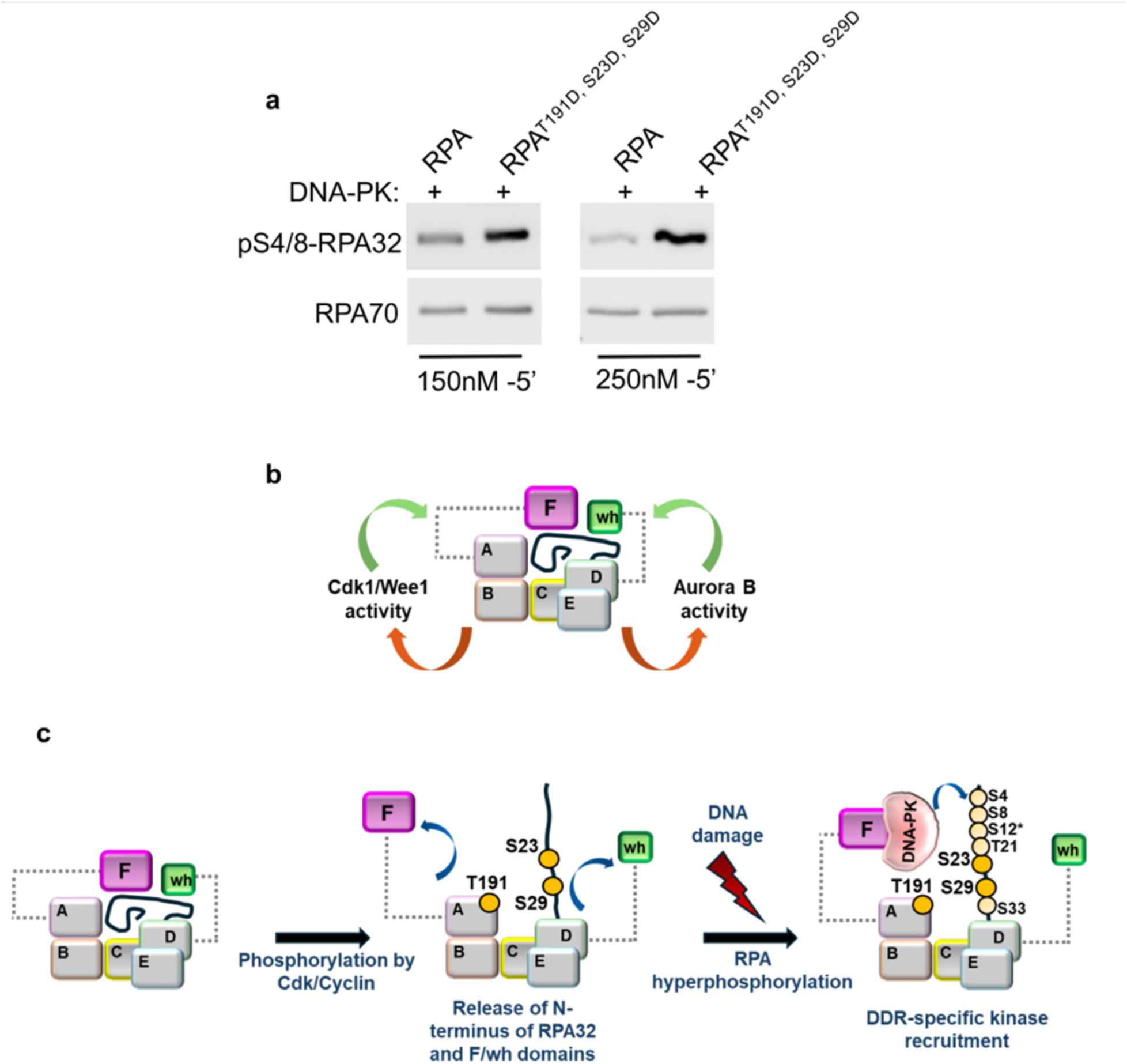
a. Cdk1 priming-driven hyperphosphorylation of RPA32. Western blot analysis of *in vitro* kinase assay of recombinant RPA (150nM or 250nM) incubated with 80 units of DNA-PK for 5 min. Blots were probed with the indicated antibodies. Data represent three independent experiments. **b.** Model depicts the positive feedback loop between RPA and the cell cycle-specific kinases that phosphorylate RPA. Domains A, B, C, D, E are depicted. It remains unclear if the activity of kinases is regulated by RPA through a direct effect on their catalytic activity or through modulation of mediators such as Wee1. The protein interaction domains (F and wh) are also shown. The disordered N-terminus of RPA32 is shown as a black line. **c.** Model illustrates the release of the N-terminus of RPA32 (black line) along with the F and wh domains upon cell cycle-specific priming phosphorylation at T191, S23, and S29 sites by Cdk1 kinase. The phosphocode primes RPA32 for efficient hyperphosphorylation by kinases such as DNA-PK in response to DNA damage. Thus, the structural re-organization induced by cell cycle-specific priming phosphorylation of both RPA70 and RPA32 work synergistically and is crucial for hyperphosphorylation of RPA32 in response to DNA damage.

## DISCUSSION

Decades ago, RPA was shown to be constitutively phosphorylated through the S and G2 to M phase of cell cycle by Cdk1 and Cdk2 kinase^13–20^. The critical roles of RPA in S-phase and DNA damage response are well-established^13^, however, RPA’s function in modulating normal cell cycle progression through the G2 and M phase has remained unclear. In addition, RPA functions have been shown to be primarily regulated through phosphorylation of the RPA32 subunit^13,14^. However, regulation of the RPA70 subunit through phosphorylation and its significance has remained obscure. We previously addressed a key function of RPA, specifically the RPA70 subunit, in dictating normal mitotic progression through regulation of Aurora B kinase activity^9^. Here we show that RPA is also important for regulating normal G2 to M transition. This solidifies a role for RPA in dictating normal cell cycle progression through G2 to M transition and for progression through mitosis.

Here, we also uncover the presence of an RPA phosphocode that spans multiple RPA subunits that can be tuned to position RPA for defined DNA metabolic functions. The spatial relationship between the protein-interaction domains of RPA and the disordered N-terminal region of RPA, and its control by phosphorylation, have remained largely unexplored^13,14^. We show that the T191 site on the RPA70 subunit also serves as a priming site for hyperphosphorylation of the RPA32 subunit in response to DNA damage, similar to the S23 and S29 sites on RPA32. Since phosphorylation at these sites are unique to normal cell cycle progression, the discovery of T191- phosphorylation presents a collective cell cycle-specific phosphocode that is critical for maintaining RPA in a configuration that is primed for DNA damage response. We also present direct evidence for such configurational changes in RPA that are induced by T191 and S23, S29 phosphorylation. These structural changes include enhanced accessibility of the F and Wh protein interaction domains of RPA, and the complete release of the N-terminal region of RPA32. Functionally, we show that one of the key outcomes of such changes is the recruitment and efficient phosphorylation of RPA by downstream kinases such as DNA-PK to mediate the genomic stress response.

Another intriguing aspect of RPA’s role in normal G2 to M progression is the feedback control of kinases that regulate these phases. Phosphorylation of the S384 residue in the RPA70 subunit by Aurora B is critical for modulating the activity of the Aurora B kinase through a positive feedback mechanism^9^. Such a positive feedback mechanism, wherein RPA regulates the activity of interacting kinases, has also been observed for ATR kinase^13,14^. Binding of ATRIP to RPA-coated ssDNA facilitates recruitment and activation of ATR^71^. However, ATR activity is not altered by S33 phosphorylation on RPA32 by ATR^13^. In this study, we show that in addition to Aurora B regulation, RPA70 is also important for maintaining Cdk1 activity through a positive feedback mechanism. This regulation is dependent on phosphorylation of RPA70 at the novel T191 site by Cdk1 (**Figure 8b**). Thus, feedback control of RPA-interacting kinases by RPA appears to be a conserved mechanism for the regulation of normal cell cycle progression and DNA damage response.

In this study, we uncover that disruption of T191 phosphorylation leads to stabilization of Wee1. However, the mechanism through which RPA regulates Wee1 remains to be resolved. It remains unclear if degradation of Wee1 kinase is regulated by a direct interaction between Wee1 and RPA. However, since Wee1 stability is also regulated by Cdk1 (through phosphorylation of Wee1) that targets it for degradation by SCFβ-TrCP, it is possible that RPA directly affects the catalytic activity of Cdk1 that in turn controls Cdk1-dependent degradation of Wee1. Future studies will parse the precise mechanism involved.

Another intriguing observation for the RPA-T191A mutant is that the inhibition of Cdk1 activity is eventually compensated, and cells are able to enter mitosis. This could be due to the effect of the Cdc25 phosphatase family that can eventually dephosphorylate Cdk1 and permit cells to enter mitosis. It is also likely that the effect of targeting the T191 site in modulating Cdk1 activity could be eventually compensated by the S23 and S29 phosphosites in G2. Expression of the S23 and S29 phosphodead mutant with knockdown of endogenous RPA showed a similar cell cycle delay in G2 phase^21^. However, the underlying mechanisms causing the delay were not characterized. It is likely that targeting the S23 and S29 phosphosites causes a similar G2 delay by inhibiting Cdk1 activity through stabilization of Wee1. Future studies will focus on the S23A/S29A, and the T191A/S23A/S29A triple mutant, to further determine the functional synergism among the phosphosites. Other posttranslational modifications such as sumoylation, acetylation, and additional phosphorylation at S135 and T180 positions have also been captured in RPA70^13^. It remains to be determined if the cell cycle-specific phosphocode is also critical for priming other such modifications.

## Supporting information

Supplementary-All

## ACKNOWLEDGEMENTS

This work was supported through grants from the National Institutes of Health, NIGMS R01GM143179 and SLU (S.O.), R35GM149320 (E.A.), and from the Office of the Director S10OD030343 (E.A.). We thank Dr. Brunda Tumala for technical support. We thank Dr. Grzegorz Sabat (University of Wisconsin Madison-Mass Spectrometry Core), and GESC@MGI at Washington University in St. Louis for their technical support.

## MATERIALS AND METHODS

### Cell culture and generation of knock-in cell line

The human colorectal carcinoma cell line (HCT116) was obtained from ATCC and maintained in McCoy’s 5A (Gibco) medium with 10% FBS (Gibco), 100 units/ml penicillin and 100 mg/ml streptomycin (Gibco). All cell lines are routinely screened for absence of mycoplasma (ATCC kit). The RPA70-T191A mutant clones were generated by the Genome Engineering & Stem Cell Center (GESC@MGI) at Washington University in St. Louis. The synthetic gRNA targeting the sequence (5′-ACTTGGACTGGTAAGGAGTG<u>NGG</u>) and the following donor single-stranded oligodeoxynucleotides (ssODN) were obtained from IDT: 5’- cccagcctgtcacacacttctgggggaacacagtccaaagtggtgcccattgccagcctcGcAccttaccagtccaagtgagttgttgcatagagtaagttcagagtgtacttatgaaatcggag). Briefly, HCT116 cells were transfected with gRNA and Cas9 complex along with ssODN. Single cell clones were sorted into 96-well plated and homozygous knock-in mutation was identified using Next Generation Sequencing. Positive clones were expanded and cryopreserved. All clones were negative for mycoplasma and authenticated by STR profiling (GESC@MGI).

### Immunoprecipitation of endogenous RPA

For immunoprecipitation (IP) of endogenous RPA70 in mitotic cells, HCT116 cells were arrested in prometaphase with nocodazole (50ng/ml) for 18 h. The mitotic cells were collected by shake-off method and washed twice in 1X phosphate buffered saline (PBS) for 2 min at 2,000 rpm at 4°C. Cell pellets were lysed using mammalian cell lysis buffer (MCLB) (50 mM Tris-Cl pH 8.0, 5 mM EDTA, 0.5% Igepal, 150 mM NaCl) supplemented with following inhibitors: 1 mM phenylmethylsufonyl fluoride (PMSF), 1 mM sodium fluoride, 10 mM β-glycerophosphate, 1 mM sodium vanadate, 2 mM DTT, 1X protease inhibitor cocktail (Sigma–Aldrich), 1X phosphatase inhibitor cocktail (Santa Cruz Biotechnology). Cells were lysed for 15 min at 4°C on a rocker and cell lysates were collected by centrifugation at 17,968 g for 10 mins at 4°C. Total protein concentrations were determined using the Bradford assay. 2.5 mg of total cell lysate in 500 μL of MCLB with inhibitors was precleared with protein A/G plus agarose beads (Santa Cruz Biotechnology) and 1μL of normal rabbit IgG (Cell Signaling, 2729) by rotating for 30 min at 4°C. Precleared cell lysates were centrifuged at 3,500 rpm for 3 min at 4°C and incubated with 20 μL protein A/G plus agarose beads and 1 μg of anti-RPA70 antibody (Cell Signaling, 2267) and rotated overnight at 4°C. Next day, unbound lysate was removed by centrifugation at 3,000 rpm for 5 min at 4°C and beads were washed thrice with cold 1X MCLB buffer. The beads were resuspended in 30 μL MCLB buffer and eluted by boiling in 1X Laemmli buffer. The eluates were subjected to SDS-PAGE electrophoresis followed by Coomassie staining. The bands corresponding to RPA70 were excised and analyzed by Mass spectrometry

### *In vitro* kinase assay

For *in vitro* kinase assays, recombinant human RPA protein (2 μM) was incubated with 2.3 μg of recombinant Cdk1/Cyclin B kinase (Millipore, Sigma) in 1X kinase reaction buffer (50 mM Tris pH 7.4, 2 mM DTT, 10mM MgCl_2_, 200 μM ATP) for 30 min at 30°C. For phosphorylation by Cdk2/Cyclin A2, recombinant human RPA protein (3 μM) was incubated with 0.1 μg of recombinant Cdk2/CyclinA2 kinase (Promega) in 1X kinase reaction buffer (40 mM Tris pH 7.5, 50 μM DTT, 20 mM MgCl_2_, 0.1 mg/ml BSA, 250 μM ATP) for 30 min at 30°C. All the kinase reactions were stopped by incubating with 2 μl of 0.4M EDTA at RT for 2 min and flash frozen prior to MS analysis. For kinase assay with DNA-PK, 150 nM of recombinant human RPA and RPA variants (RPA^T191D^ and RPA ^T191D, S23D, S29D^) were incubated with 80 units of DNA-PK (Promega) and 10 μg/ml linear double-stranded DNA in 1X kinase reaction buffer (40 mM Tris pH7.5, 20 mM MgCl_2_, 0.1 mg/ml BSA, 1 mM DTT, 200 μM ATP) for 5 min at 30°C. Reactions were stopped by 2μl of 0.4M EDTA and by boiling in 1X Laemmli buffer followed by western blot analysis.

### Mass Spectrometry

*Enzymatic “In Liquid” Digestion: In vitro* phosphorylation assay reaction (128ul) was diluted for protein precipitation with 192ul miliQ water, 80ul saturated TCA and 400ul of acetone to final 800ul volume and sample was incubated on ice for 30 minutes prior to 10-minute centrifugation at 16,100xg. Generated pellet was washed twice in cold acetone, air-dried briefly and re-solubilized in 15μl of 8M Urea / 50mM NH_4_HCO_3_ (pH8.5) for the reduction step with: 2.5μl of 25mM DTT, and 42.5μl 25mM NH_4_HCO_3_ (pH8.5). Incubated at 56°C for 15 minutes, cooled on ice to room temperature then 3μl of 55mM CAA (chloroacetamide) was added for alkylation and samples were incubated in darkness at room temperature for 15 minutes. Reaction was quenched by adding 8μl of 25mM DTT. Finally, 10μl of Trypsin/LysC solution [100ng/μl 1:1 *Trypsin* (Promega) and *LysC* (FujiFilm) mix in 25mM NH_4_HCO_3_] and 19μl of 25mM NH_4_HCO_3_ (pH8.5) was added to 100µl final volume. Digestion was conducted o/n at 37°C. Immunoprecipitated samples were analyzed similar to the *in vitro* kinase samples as detailed above. Reaction was terminated by acidification with 2.5% TFA [Trifluoroacetic Acid] to 0.3% final.

*NanoLC-MS/MS:*Digests were desalted using Pierce™ C18 SPE pipette tips (100µl volume) per manufacturer protocol and eluted in 20µl of 70/30/0.1% ACN/H_2_O/TFA. Dried to completion in the speed-vac and finally reconstituted in 25µl of 0.1% formic acid. Peptides were analyzed by nanoLC-MS/MS using the Agilent 1100 nanoflow system (Agilent) connected to hybrid linear ion trap-orbitrap mass spectrometer (LTQ-Orbitrap Elite™, Thermo Fisher Scientific) equipped with an EASY-Spray™ electrospray source (held at constant 35°C). Chromatography of peptides prior to mass spectral analysis was accomplished using capillary emitter column (PepMap® C18, 3µM, 100Å, 150×0.075mm, Thermo Fisher Scientific) onto which 1.5µl of extracted peptides was automatically loaded. NanoHPLC system delivered solvents A: 0.1% (v/v) formic acid, and B: 99.9% (v/v) acetonitrile, 0.1% (v/v) formic acid at 0.50 µL/min to load the peptides (over a 30 minute period) and 0.3µl/min to elute peptides directly into the nano-electrospray with gradual gradient from 0% (v/v) B to 30% (v/v) B over 80 minutes and concluded with 5 minute fast gradient from 30% (v/v) B to 50% (v/v) B at which time a 5 minute flash-out from 50-95% (v/v) B took place. As peptides eluted from the HPLC-column/electrospray source survey MS scans were acquired in the Orbitrap with a resolution of 120,000 followed by CID-type MS/MS fragmentation of 30 most intense peptides detected in the MS1 scan from 350 to 1800 m/z; redundancy was limited by dynamic exclusion.

*Data analysis:* Elite acquired raw MS/MS data files were converted to mgf file format using MSConvert (ProteoWizard: Open-Source Software for Rapid Proteomics Tools Development). Resulting mgf files were used to search against *Homo sapiens* UP000005640 reference proteome database (81,838 total entries; 02/01/2023 download) along with a cRAP common lab contaminant database (116 total entries) using in-house Mascot search engine 2.7.0 [Matrix Science] with fixed Cysteine carbamidomethylation and variable Serine and Threonine phosphorylation, Methionine oxidation plus Asparagine or Glutamine deamidation. Peptide mass tolerance was set at 10ppm and fragment mass at 0.6 Da. Protein annotations, significance of identification and spectral based quantification was done with Scaffold software (version 5.0.1, Proteome Software Inc., Portland, OR). Peptide identifications were accepted if they could be established at greater than 60.0% probability to achieve an FDR less than 1.0% by the Scaffold Local FDR algorithm. Protein identifications were accepted if they could be established at greater than 5.0% probability to achieve an FDR less than 1.0% and contained at least 2 identified peptides. Protein probabilities were assigned by the Protein Prophet algorithm (Nesvizhskii, Al et al Anal. Chem. 2003;75(17):4646-58). Proteins that contained similar peptides and could not be differentiated based on MS/MS analysis alone were grouped to satisfy the principles of parsimony. Proteins sharing significant peptide evidence were grouped into clusters.

### Western blot analysis

For western blotting, cells were lysed in MCLB buffer as described above. Proteins were resolved by SDS-PAGE and transferred on to nitrocellulose membrane (0.45um; Bio-Rad laboratories). Blots were blocked in 5% non-fat dry milk dissolved in Tris-Buffered Saline with 0.1% Tween-20 (TBS-T) at RT for 1 h. The following antibodies were diluted in TBS-T buffer and incubated overnight at 4°C, except β-Actin which was incubated at room temperature for 30 min: β-Tubulin (1:2000; Cell Signaling, 2128), Vinculin (1:2000; Cell Signaling, 13901), β-Actin (1:1000; Cell Signaling, 3700), RPA70 (1:2000; Cell Signaling, 2198), p53 (1:1000; Santa Cruz Biotechnology, sc-126), RPA32 (1:5000; Cell Signaling, 35869), Cdk1 (1:1000; Santa Cruz Biotechnology, sc-54), Wee1 kinase (1:1000; Cell Signaling, 13084), Cyclin B1 (1:1000; Santa Cruz Biotechnology, sc-24). The following phospho-specific antibodies diluted in TBS-T with 1% milk were incubated overnight at 4°C: pS384-RPA70 (custom antibody synthesized by Genscript described previously^9^), pS10-Histone H3 (Cell Signaling, 53348), pS4/S8-RPA32 (1:2000; Bethyl, A-300-245A), pS317-Chk1 (1:1000, Cell Signaling,), pS345-Chk1 (1:1000; Cell Signaling, 2341), S15-p53 (1:1000; Cell Signaling, 9284) or diluted with 5% milk: pY15-Cdk1 (1:1000; Santa Cruz Biotechnology, sc136014) and pS33-RPA32 (1:1000; Bethyl, A300-246A). Membranes were probed with HRP-conjugated secondary antibodies 1:30,000 diluted in TBS-T buffer (Jackson Immunoresearch). Blots were developed with ECL substrate (Pierce) and images were acquired in the iBright CL1500 imager (Thermo Fisher Scientific). Blots were quantitated using iBright analysis software.

### Viability assays

Cell proliferation rate was assessed using CellTiter 96 Aqueous One Solution Cell Proliferation (MTS) assay (Promega). Briefly, 5 x10^3^ HCT116 cells were seeded in 96-well plate and 20 μL of MTS reagent was added to each well. The plates were incubated at 37°C for 1 h and absorbances were recorded at 495nm (Synergy H1-BioTek). To calculate cell proliferation rate, all absorbances were normalized to absorbance at 0 h for respective samples. Apoptosis was measured using Caspase-Glo® 3/7 assay (Promega). Briefly, 10 x 10^3^ cells were cultured in 96-well microplates (Corning) and treated with either 100 ng/ml SN-38 (Tocris) or vehicle (0.1% DMSO) for 21 h. Cells were incubated with 100 μL Caspase-Glo reagent for 90 min at room temperature (RT) in dark and plates were scanned using a luminescence plate reader (Synergy H1-BioTek). Background-corrected values were obtained by subtracting blank (media and reagent) from other samples.

### Cell synchronization

Cells were synchronized at G1/S border using double thymidine arrest. Briefly, 1 × 10^6^ *RPA^WT^* cells and 1.2 × 10^6^ RPA*^T191A^* cells were seeded per 10 cm dish and next day, once cells reached 50-60% confluency, they were treated with 2 mM thymidine (Sigma-Aldrich) for 16 h. Cells were washed twice with 1X PBS and cultured in fresh McCoy’s medium for 8 h. For the second thymidine arrest, cells were cultured in media supplemented with 2mM thymidine for 16 h. Cells were then released from arrest by washing twice with 1X PBS and cultured in fresh media without drug. For 0 h analysis, cells were collected just prior to release. After release from the G1/S border, cells were harvested at 3 h, 6 h for flow cytometric analysis and at 7 h and 8.5 h for western blotting. Cells released into G2 phase at 7 h were treated with 100 μM Etoposide (Sigma) or 0.1% DMSO (vehicle control) for 45 min. Cells were lysed in MCLB buffer as described above. HCT116 cells were synchronized in mitosis (pro-metaphase) using 75 ng/mL nocodazole (Sigma) for 18 h and mitotic cells were collected by shake off method and lysed as indicated above.

### Flow cytometry

For flow cytometric analysis, cells were trypsinized for 5 min at 37°C and resuspended in McCoy’s 5A medium. Cells were centrifuged at 931*g* for 2 min at 4°C and the pellet was washed once with 1X PBS. Cells were resuspended in 500 μL of 1X PBS and added dropwise to 5 mL of 100% ethanol and re-suspended by mildly vortexing the samples. For propidium iodide staining, ethanol-fixed samples were centrifuged at 2,000 rpm at 4°C for 2 min and cells were resuspended in 1% BSA dissolved in 1X PBS. For propidium iodide (PI) staining, cells were counted and 1×10^6^ cells were resuspended in 3/50 volume of 50X Propidium Iodide solution, 1/40 volume of 10 mg/mL RNase A (Thermo Scientific) and 1% BSA dissolved in 1X PBS. Samples were filtered through 35 μm cell stainer cap and collected in polystyrene tubes (Corning). DNA content was analyzed by BD FACS Canto II flow cytometer (BD Biosciences) and the samples were collected using a low flow rate, and a minimum of 15,000 cycling events were recorded. Cell cycle distribution was analyzed using ModFit LT (verity) software.

### Immunofluorescence

HCT116 cells were cultured on 12 mm glass coverslips (Neuvitro) in 12-well culture plates. Cells were washed once with 1X PBS and fixed in 4% paraformaldehyde overnight at 4°C. Next day, coverslips were rinsed three times with 1X PBS and permeabilized with 2% Triton-X100 in PBS for 15 min, followed by 30 min incubation in blocking solution (2% BSA, 0.1%Igepal, 1X PBS). For immunostaining, all antibodies were diluted in blocking solution and the following primary antibodies were used; α-Tubulin (1:45; Cell Signaling, 2125), pS10-Histone H3 (1:1000; Millipore Sigma, 05-1336). Cells were incubated with primary antibody for 1h at RT and coverslips were washed three times in wash buffer (0.1% Igepal in 1X PBS). The following secondary antibodies were used: goat anti-rabbit Alexa Fluor 488 (1:500; Thermo Fisher) and donkey anti-mouse Alexa fluor 545 (1:500; Thermo Fisher) and incubated for 45 min at RT.

Coverslips were mounted on glass slides (Fisher) and mounted with ProLong Gold antifade reagent with DAPI (Invitrogen). Samples were analyzed with Leica DM6 B upright fluorescent microscope with a 100X oil immersion objective and images were taken with Leica DFC 9000GT camera. Images were processed using LAS X software with linear LUT covering full data range. To determine the percentage of mitotic cells, pS10 Histone H3 stained cells were analyzed using ImageJ software. Mitotic cells undergoing anaphase and telophase were counted as one cell. The total number of cells per image stained blue (DAPI) and pink (pHH3) were counted using ImageJ. The images were converted from an RGB to an 8-bit image followed by background subtraction. Images were subjected to watershed processing that separated nuclei from each other followed by counting of nuclei. This overlay was then checked manually for non-specific counts that were subtracted from the total count.

### Purification of RPA and phosphomimetic variants

Human RPA was produced using plasmid pET-Duet1-hRPA^syn^-70C-His coding for a poly-His affinity tag at the C-terminus of RPA70. The open reading frames for RPA70, RPA32, and RPA14 were codon-optimized for overexpression in *E. coli* (GenScript Inc). RPA70 was engineered into multiple cloning site (MCS) 1 while RPA32 and RPA14 were cloned into MCS2. RPA was purified as described^9^ with the following modifications. Point mutations coding for the T191D, S23D, and or S29D substitutions were generated by GenScript Inc. Briefly, the respective plasmid was transformed into BL21 (DE3) cells and transformants were selected using ampicillin (100 μg/mL). A single colony was inoculated in 1 L of Luria Broth and incubated at 37 °C without shaking for 20-24 hrs. Cells were then grown at 37 °C with shaking at 250 rpm, until the OD_600_ reached 0.6 and then induced with 0.4 mM Isopropyl b-D-1-thiogalactopyranoside (IPTG). Induction was carried out at 37 °C for 3 hrs. Harvested cells were resuspended in 30 mL/L cell resuspension buffer (30 mM HEPES, pH 7.8, 300 mM KCl, 0.02% Tween-20, 1.5 X protease inhibitor cocktail, 1 mM PMSF, and 10% (v/v) glycerol). Cells were lysed with 0.4 mg/mL lysozyme for 30 min at 4 °C followed by sonication in the cold room. The samples were maintained at 4 °C through the entirety of the remaining purification. The clarified lysate was fractionated over a Ni^2+^-NTA agarose column (Gold Biotechnology Inc.). After loading, the column was first washed with resuspension buffer containing 2M NaCl to remove any trace non-specifically bound nucleic acids. A subsequent wash with resuspension buffer was performed to equilibrate the sample to buffer containing 300 mM KCl. RPA was eluted using cell resuspension buffer containing 400 mM imidazole. Fractions containing RPA were pooled and diluted with H_0_ buffer (30 mM HEPES, pH 7.8, 0.02% Tween-20, 1.5X protease inhibitor cocktail, 10% (v/v) glycerol and 0.25 mM EDTA pH 8.0) to match the conductivity of buffer H_100_ (H_0_ + 100 mM KCl), and further fractionated over a fast-flow Heparin column (Cytiva Inc.). RPA was eluted using a linear gradient of H_100_–H_1500_ buffers (subscripts denote the mM concentrations of KCl), and fractions containing RPA were pooled and concentrated using an Amicon spin concentrator (30 kDa molecular weight cut-off). The concentrated RPA was further fractionated over a HiLoad 26/600 Superdex-200 column (Cytiva Inc.) using RPA storage buffer (30 mM HEPES, pH 7.8, 200 mM KCl, 0.25 mM EDTA, 0.01% Tween-20, and 10% (v/v) glycerol). Purified RPA was flash frozen using liquid nitrogen and stored at –80°C. RPA concentration was measured spectroscopically using ε_280_ = 87,410 M^−1^cm^−1^. The phosphomimetic RPA variant proteins were also purified using the same procedure.

### Circular Dichroism measurements of secondary structure

CD measurements were performed using a Chirascan V100 spectrometer (Applied Photophysics Inc.). A nitrogen fused set up with a cell path of 0.5 mm was used to perform the experiments at 20 °C. All CD traces were obtained between 200-260 nm, and traces were background corrected using CD reaction buffer (100 mM NaF, 1 mM TCEP-HCl, and 5 mM Tris-Cl, pH 7.5). 3 µM of RPA or RPA phosphomimetic variants were used, and 10 scans were collected and averaged per sample using 1 nm step size and 1 nm bandwidth.

### RPA-ssDNA binding measured using fluorescence anisotropy

5’-FAM-(dT)_20_ or 5’-FAM-(dT)_35_ were diluted to 10 nM in DNA binding buffer (50 mM Tris acetate pH 7.5, 50 mM KCl, 5 mM MgCl_2_, 1 mM DTT, 10 % (v/v) glycerol, and 0.2 mg/ml BSA). 1.2 ml of this ssDNA stock was taken in 10 mm pathlength quartz cuvettes (Firefly Inc.) maintained at 23 °C and G-factor corrected fluorescence anisotropy was measured (in triplicate) using a PC1 spectrofluorometer (ISS Inc.). Data acquisition was performed using the instrument-associate Vinci 3 software. Samples were excited at 488 nm and the resulting fluorescence emission was collected using a 520 nm bandpass emission filter. To extract corrected anisotropy from raw values (instrument readings) the concentrations of ssDNA, total fluorescence, and the added protein concentration were first corrected for the effect of sample dilution resulting from the stepwise addition of the protein. Second, the FAM anisotropy was corrected for changes in the fluorescence quantum yield of the bound species due to proximity of the fluorescein moiety to the protein in order to plot the anisotropy. For this, the dilution-corrected FAM fluorescence values were used to correct and rescale anisotropy values as described^9^. The saturation points were taken as the intersection of biphasic curves from the linear fits of the initial data points reflecting the change in anisotropy upon binding of sub-saturating amounts of proteins.

### Measurement of ssDNA binding kinetics using stopped-flow fluorescence

Stopped flow experiments were performed on a SX20 instrument (Applied Photophysics Inc.) at 25 °C in RPA reaction buffer (30 mM HEPES pH 7.8, 100 mM KCl, 5 mM MgCl_2_, 6 % (v/v) glycerol, and 1 mM β-mercaptoethanol). Seven to eight individual shots were averaged for each experiment. All experiments were repeated a minimum of n=3 times and the mean value and SEM calculated from the individual fits are reported. RPA binding to ssDNA was captured by monitoring the change in intrinsic Trp fluorescence. RPA (200 nM) from one syringe was mixed with increasing concentrations of (dT)_35_ ssDNA (0-400 nM) from the other syringe and the change in Trp fluorescence was measured as a function of time by exciting the sample at 290 nm and collecting fluorescence emission using a 305 nm long-pass filter. Data were fit to a single exponential equation and a plot of the k_obs_ (s^-1^) as a function of [RPA] yielded k_on_ and k_off_ values.

### Mass photometry (MP) measurements

All MP measurements were carried out on a TwoMP instrument (Refeyn Ltd.) as described^10^. Briefly, glass coverslips (No. 1.5H thickness, 24 × 50 mm, VWR) were cleaned by sonication in isopropanol followed by deionized water and dried using a nitrogen gas stream. For each round of measurement, a clean coverslip was placed on the oil-immersion objective lens (Olympus PlanApo N, 1.42 NA, 60×), with a holey-silicone gasket (Refeyn Ltd) adhered on the top surface of the coverslip. All dilutions and measurements were performed at room temperature (23 ± 2°C) in 1× buffer (20 mM HEPES, pH 7.5 and 150 mM KCl) supplemented with 1 mM DTT. Samples of DNA alone, RPA alone, or ssDNA-RPA mixtures were allowed to equilibrate at 23 ± 2°C for 5 min after which 1 μl of the respective sample was quickly diluted in 15 µL of buffer. The newly adhered spots were video recorded for 1 min. High contrast (light-scattering) events corresponding to single particle landings on the coverslip were analyzed further. A known mass standard (β-amylase, Sigma A8781-1VL) was used to convert image contrast-signal into mass units. Histograms were plotted from all the data gathered during the 1 min video interval and non-linear least squares fit to single gaussian function to extract mean mass and error.

### Crosslinking Mass Spectrometry (XL-MS) analysis

RPA or the phosphomimetic variants (10 μM) were incubated with 5 mM BS3 crosslinker (Millipore Sigma Inc.) at room temperature for 15 min in 20 μl reaction buffer (50 mM HEPES pH 7.8, 100 mM KCl, 10% glycerol). The crosslinking reaction was quenched with 2 μl of 1M ammonium acetate for 15 min and the samples were separated on SDS-PAGE. Gel bands were excised and destained with a 50 mM ammonium bicarbonate and 50% acetonitrile mixture and reduced with a mixture of 100 mM dithiothreitol and 25 mM bicarbonate for 30 minutes at 56 °C. The reaction was subsequently exchanged for the alkylation step with 55 mM iodoacetamide and 25 mM ammonium bicarbonate and incubated in the dark at room temperature for 25 min. The solution was then washed with the 50 mM ammonium bicarbonate and 50 % acetonitrile mixture. The gel pieces were then first dehydrated with 100 % acetonitrile and then rehydrated with sequence grade trypsin solution (0.6 µg, Promega) and incubated overnight at 37 °C. The reaction was quenched with 10 µL of 50% acetonitrile and 0.1% formic acid (FA, Sigma) and transferred to new microfuge tubes, vortexed for 5 min, and centrifuged at 15,000 rpm for 30 min. The extracted and dried peptide was reconstituted with 0.1% FA in water and injected onto a Neo trap cartridge coupled with an analytical column (75 µm ID x 50 cm PepMap Neo C18, 2 µm). Samples were separated using a linear gradient of solvent A (0.1% formic acid in water) and solvent B (0.1% formic acid in ACN) over 120 mins using a Vanquish Neo UHPLC System coupled to an Orbitrap Eclipse Tribrid Mass Spectrometer with FAIMS Pro Duo interface (Thermo Fisher Scientific). The acquired MS/MS data was queried for cross-link identification against the sequence of 5 target proteins using Proteome Discoverer v3.0 with the XlinkX node, applying a 1% FDR for cross-link validation. Data were visualized using xiVIEW and plotted in Inkscape.

### Size exclusion chromatography analysis of RPA-ssDNA complexes

SEC experiments to capture formation of RPA nucleoprotein filaments on ssDNA were performed using 600 μl of 3 μM RPA in the absence or presence of ssDNA 1 µM (dT)_97_. RPA and ssDNA were premixed and incubated for 10 min on ice and resolved using a Superose 6 Increase (10/300) size exclusion column (Cytiva) equilibrated with SEC buffer (30 mM HEPES, pH 7.8, 100 mM KCl, 1 mM TCEP-HCl, and 10 % (v/v) glycerol). 0.5 ml fractions were collected and assessed using SDS-PAGE.

