## Supplementary-All for "Mechanism of RPA phosphocode priming and tuning by Cdk1/Wee1 signaling circuit"

Supplementary Figures 1-5

**Figure S1.**

**a** Peptide: VVPIASLTPYQSK

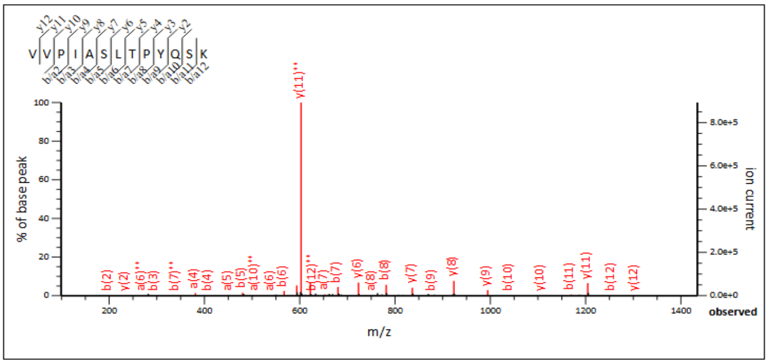


**b**

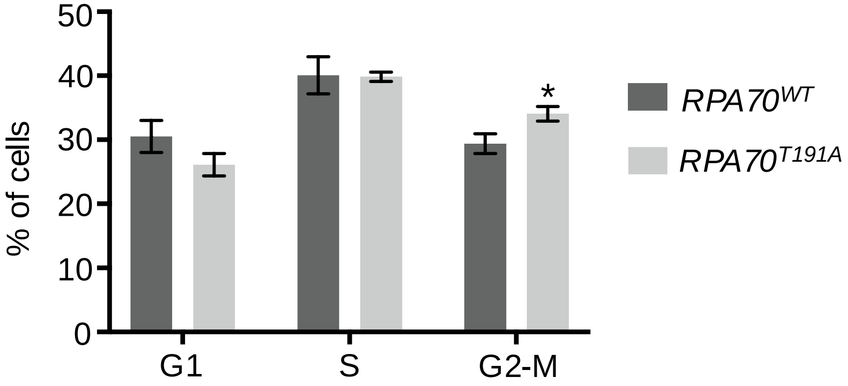


**Supplementary Figure S1. RPA70-T191 phosphorylation is not mediated by Cdk2 and increased G2/M population in the phosphodead mutant. a.** Mass spectrum of the RPA70 tryptic peptide extracted from *in vitro* kinase assay of recombinant human RPA incubated with Cdk2/Cyclin A2 complex, followed by MS-MS analysis. 16 independent MS/MS spectra of the indicated peptide observed but no phosphorylation sites were detected. **b.** Cell cycle profile of asynchronous cells analyzed by flow cytometry and DNA content assessed by propidium iodide staining. Bar graph shows mean of three independent experiments. Statistical significance determined using an unpaired one-tailed *t*-test: *$p$=0.03. Error = SEM.

**Figure S2.**

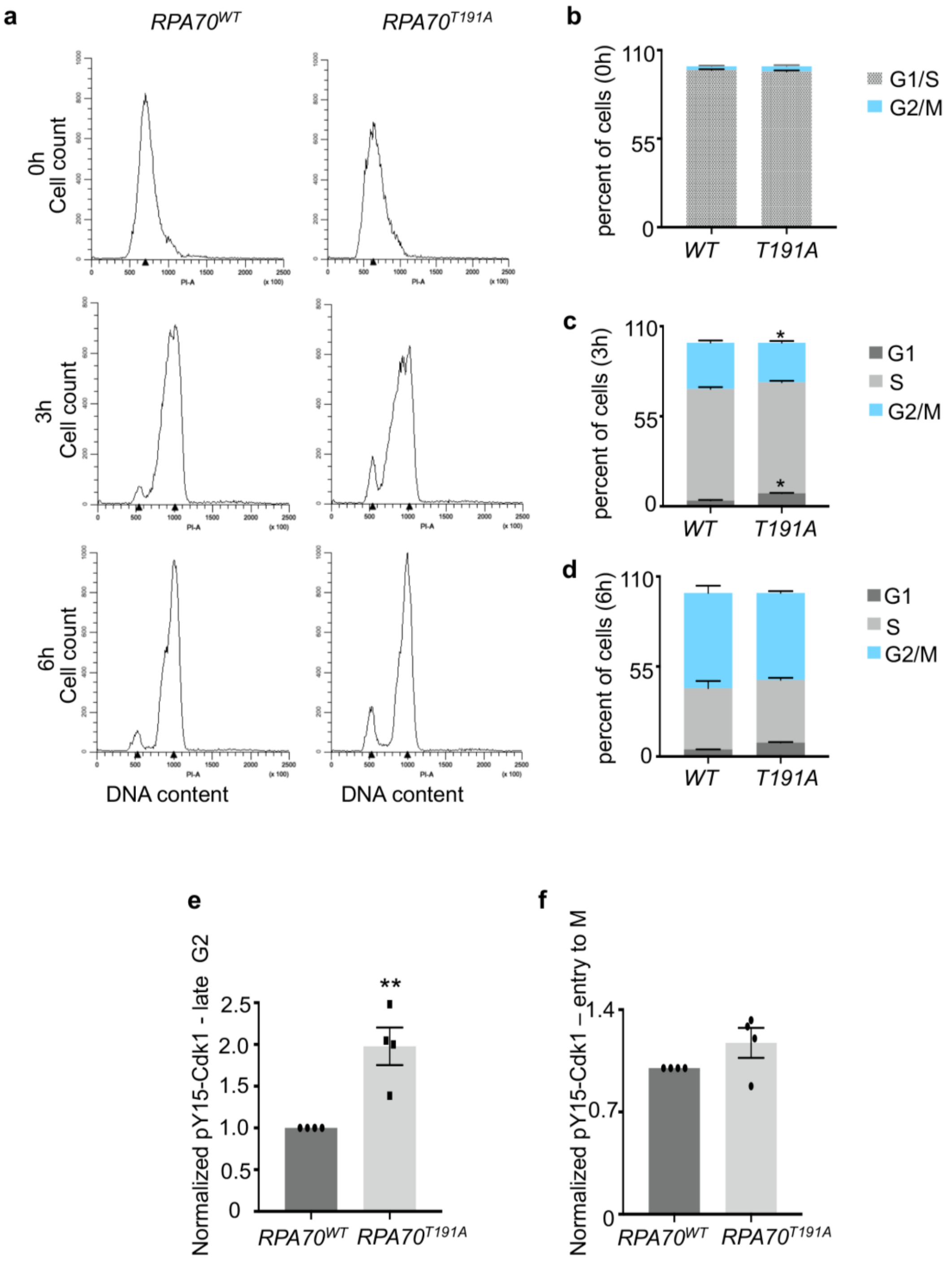


**Supplementary Figure S2. Synchronization by double thymidine block and release shows increased phospho-Y15-Cdk1 levels in late G2 of mutant cells. a.** Cell cycle profiles of cells synchronized by double thymidine block and analyzed by flow cytometry. DNA content determined using propidium iodide staining. Data represent three independent experiments. Cells arrested at 0 hrs were released and analyzed at 3 hrs and 6 hrs post-release. **b, c and d.** Profiles shown in **a.** were quantitated and the number of cells per cell cycle phase were assessed. Since cells were arrested at the G1/S border, percent cell cycle distribution at 0 hrs depicts G1 and S phase combined. Error=SEM. Statistically significant differences determined using a two-way ANOVA, Sidak's multiple comparisons test. No significant differences observed except for the mild difference for G1 and G2/M phase at 3hrs indicated by *p=0.017 and *p= 0.028, respectively. **e and f.** Western blots represented in Figure 3a, were quantitated and the levels of phospho-Y15-Cdk1 determined at 7 hrs (G2 phase) (e) and 8.5 hrs (entry to M) (f) post-release from double thymidine block. Bar graphs represent mean of three independent experiments. Error=SEM. Statistical significance determined using an unpaired two-tailed t-test with $**p = 0.0049$ (**e**) and $p = 0.1413$ (ns) (**f**).

**Figure S3.**

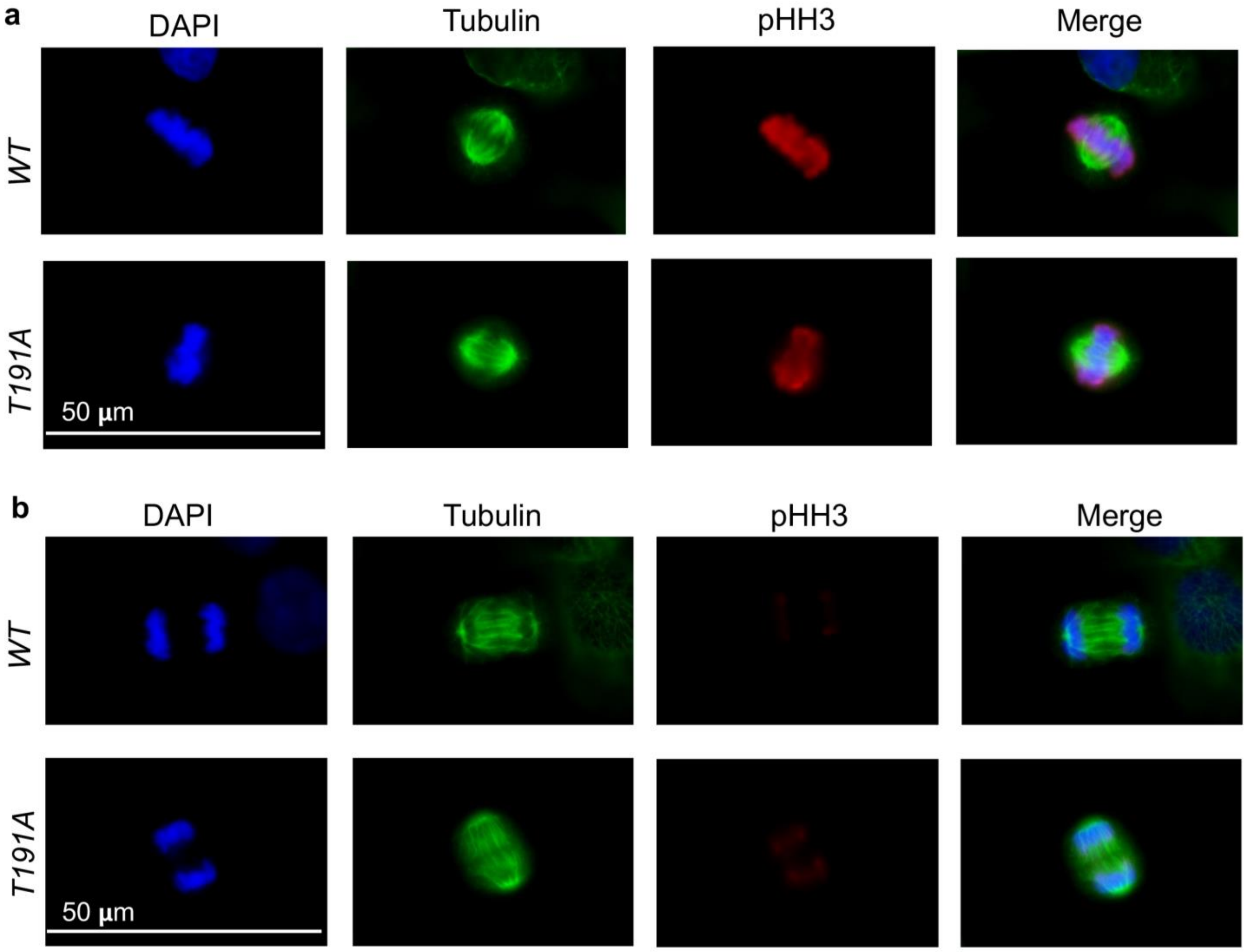


**Supplementary Figure S3. Loss of T191 phosphorylation does not affect progression through mitosis. a.** and **b.** Representative immunofluorescent images stained with DAPI, anti-Tubulin antibody and anti-phospho-S10-Histone H3 antibody and their overlay are shown. Cells in metaphase (**a**) or anaphase (**b**) in an asynchronous population are shown. No overt changes in chromosome segregation, chromosome condensation or spindle assembly observed. Data represent three independent experiments.

**Figure S4.**

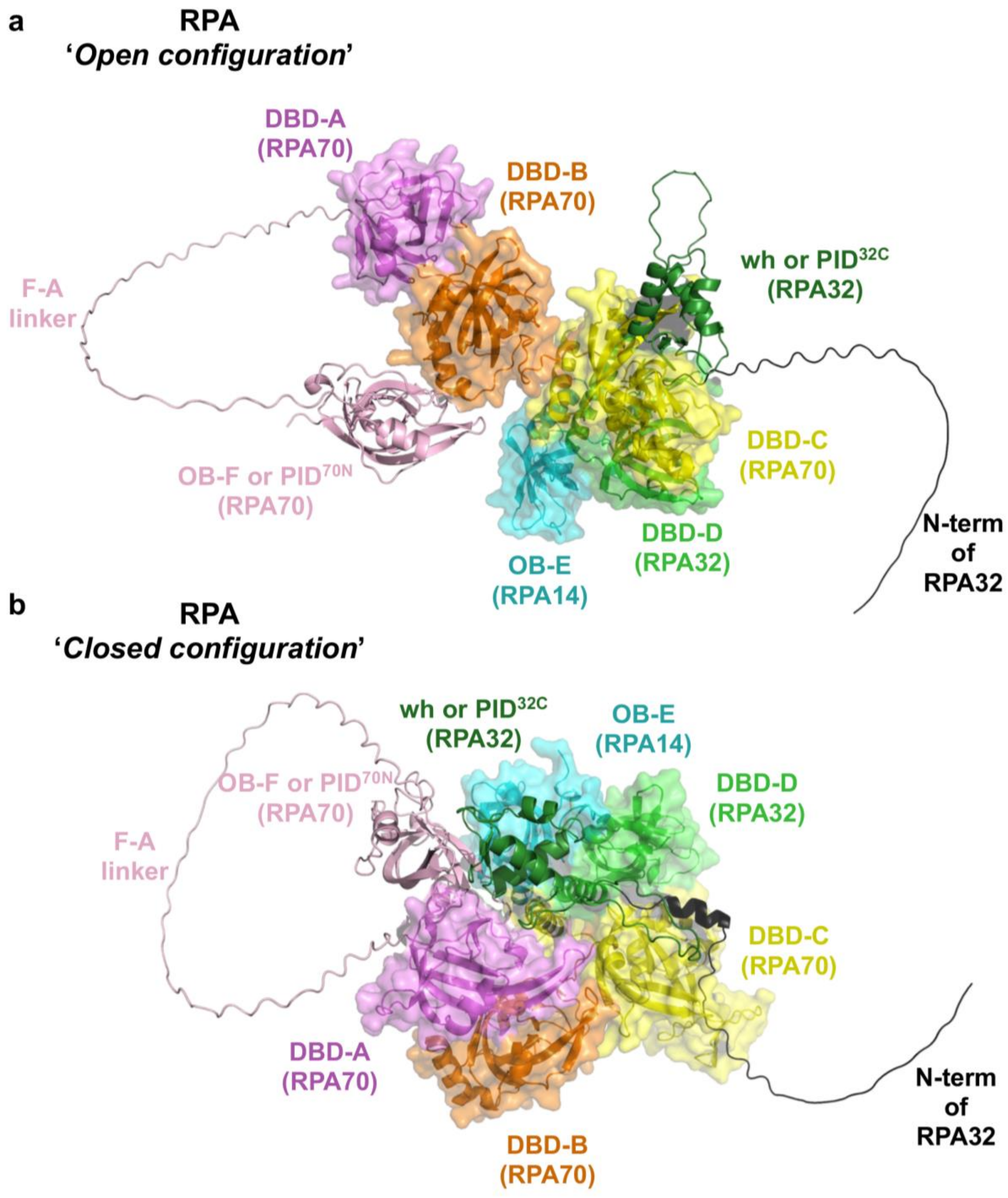


**Supplementary Figure S4. Models of the open and closed forms of RPA. a and b.** AlphaFold3 models of RPA shows **a**) an open versus **b**) closed configuration of the various domains. The individual DNA binding and protein interaction domains are shown along with their respective disordered linkers. The N-terminus of RPA32 (which harbors the sites of hyperphosphorylation) is depicted in black. In the predicted open form of RPA, the $PID^{70N}$ and $PID^{32C}$ protein interaction domains are not in proximity (panel a). These domains are together in the closed configuration (panel b).

**Figure S5.**

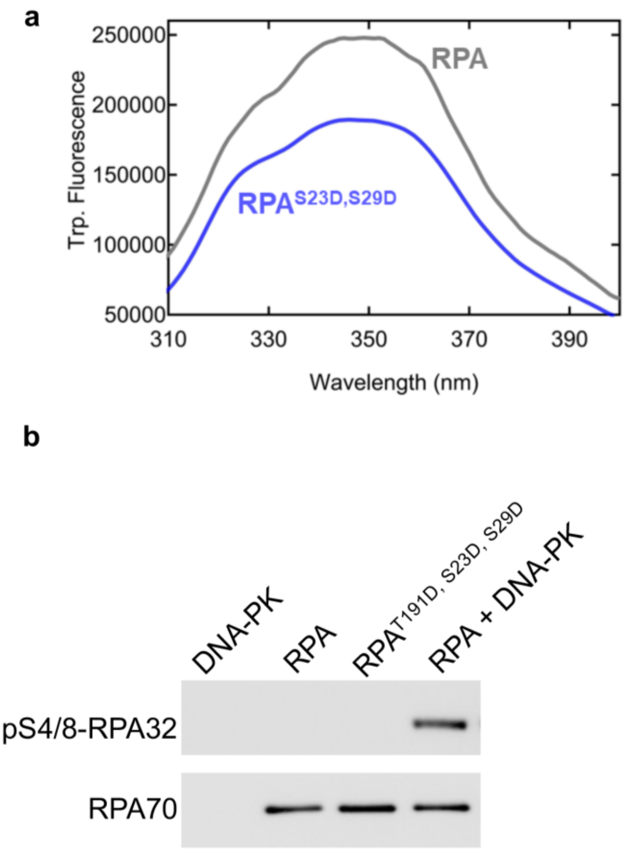


**Supplementary Figure S5. a.** Intrinsic Trp fluorescence scan of RPA versus RPA-S23D, S29 shows reduced signal for the phosphomimetic mutant. **b.** Western blot analysis of *in vitro* kinase assay of DNA-PK only, recombinant RPA only or RPA (150nM) incubated with DNA-PK for 5 min. Blots were probed with the indicated antibodies. Data represents three independent experiments.
